# Expression levels of long noncoding natural antisense transcripts overlapping the *UGT73C6* gene affect rosette size of *Arabidopsis thaliana*

**DOI:** 10.1101/2021.03.01.433436

**Authors:** Shiv Kumar Meena, Michel Heidecker, Susanne Engelmann, Ammar Jaber, Tebbe de Vries, Katja Baumann-Kaschig, Steffen Abel, Sven-Erik Behrens, Selma Gago-Zachert

## Abstract

Natural antisense long noncoding RNAs (lncNATs) are involved in the regulation of gene expression in plants, modulating different relevant developmental processes and responses to various stimuli. We identified and characterized two lncNATs (*NAT1_UGT73C6_* and *NAT2_UGT73C6_*, collectively *NATs_UGT73C6_*) in *Arabidopsis thaliana* that are transcribed from a gene overlapping *UGT73C6*, a member of the *UGT73C* subfamily of genes encoding UDP-glycosyltransferases (UGTs). Expression of both *NATs_UGT73C6_* is developmentally controlled and occurs independently of the transcription of *UGT73C6* in *cis*. Downregulation of *NATs_UGT73C6_* levels through artificial microRNAs results in a reduction of the rosette area, while constitutive overexpression of *NAT1_UGT73C6_* or *NAT2_UGT73C6_* leads to the opposite phenotype, an increase in rosette size. This activity of *NATs_UGT73C6_* relies on its RNA sequence, and, although modulation of *UGT73C6* in *cis* cannot be excluded, the observed phenotypes are not a consequence of the regulation of *UGT73C6* in *trans*. The *NATs_UGT73C6_* levels were shown to affect cell proliferation and thus individual leaf size. Consistent with this concept, our data suggest that the *NATs_UGT73C6_* modulate the expression levels of key transcription factors involved in regulating leaf growth by modulating cell proliferation. These findings thus reveal an additional regulatory layer on the process of leaf growth.

## Introduction

Long noncoding RNAs (lncRNAs) are transcripts of more than 200 nucleotides without protein-coding capacity (Mercer et al., 2009). Many lncRNAs have been shown to be important regulators of gene expression (Rinn and Chang, 2012). lncRNAs exert their activity in diverse cellular contexts and participate in different biological processes regulating chromatin organization and transcription, mRNA stability and translation (Yao et al., 2019). Based on their relative location to nearby protein-coding genes lncRNAs are classified as long noncoding natural antisense transcripts (lncNATs) and intronic, intergenic or promoter lncRNAs (Rinn and Chang, 2012; Ariel et al., 2015). lncNATs are transcripts generated from the DNA strand opposite to a protein-coding gene and overlap at least one coding exon (Ariel et al., 2015). Natural antisense transcripts (NATs) are distinguished into *cis*- and *trans-*NATs. In the case of *cis*-NATs, sense and antisense transcripts are generated from the same genomic locus and show perfect complementarity, while *trans*-NATs are transcribed from a different locus and share only partial sequence complementarity with the sense transcript (Lapidot and Pilpel, 2006). The mode of action of lncNATs to regulate their targets is diverse. Thus, in response to certain environmental or developmental stimuli, gene expression has been shown to be post-transcriptionally downregulated by endogenous, small interfering RNAs (nat-siRNAs), which are generated by RNAi from hybrids from sense and antisense RNAs (Borsani et al., 2005; Zhang et al., 2013). Additionally, it was proposed that lncNATs can act in *trans*, particularly when they are co-expressed and share high levels of sequence similarity with their potential targets (Wang et al., 2006). This could enable lncNATs to control the expression of more than one gene in the context of multigene families. lncNATs were further shown to regulate gene expression through transcriptional collision, DNA methylation, histone modification, alteration of mRNA stability, promotion of endogenous siRNA formation and by modulation of mRNAs splicing, editing and translation (Lin et al., 2015).

lncRNAs have been largely studied in mammalian and, particularly in human cells. Less is known about their function in plants, where only a few lncNATs have been functionally characterized (Wang and Chekanova, 2017). These lncNATs are important regulators of gene expression and are involved in the modulation of developmental processes and in the responses to different stimuli (Yu et al., 2019). In the model plant *Arabidopsis thaliana*, lncNATs are part of the control circuits of fundamental processes including germination (Fedak et al., 2016), flowering (Csorba et al., 2014; Rosa et al., 2016; Henriques et al., 2017; Zhao et al., 2018) and gametophyte development (Wunderlich et al., 2014). Furthermore, it has been reported that an antisense lncRNA is involved in cold acclimation (Kindgren et al., 2018). In rice, lncNATs modulate responses to phosphate starvation (Jabnoune et al., 2013), and are involved in the maintenance of leaf blade flattening (Liu et al., 2018). In tomato, a particular lncNAT, when overexpressed, was shown to confer resistance to the oomycete *Phytophthora infestans* (Cui et al., 2017).

Plant uridine diphosphate (UDP) glycosyl transferases (UGTs) are enzymes that transfer UDP-activated sugars to a variety of aglycone substrates, including hormones, secondary metabolites and xenobiotics (Ross et al., 2001). Glycosylation leads to changes in the target properties that alter their bioactivity and solubility, and plays an important role in maintaining cellular homeostasis by regulating the activity and location of important cellular metabolites and hormones (Li et al., 2001). In *A. thaliana*, the UGT family contains 107 putative genes and 10 pseudogenes organized in 14 distinct groups named A to N (Ross et al., 2001; Caputi et al., 2012). The *UGT73C* subfamily, which is part of group D, consists of seven genes. The *UGT73C1* to *UGT73C6* genes are clustered in a tandem repeat on chromosome 2 and exhibit a high nucleotide identity (between 77% and 91%). The *UGT73C7* gene is located on chromosome 3 and shows the lowest levels of similarity (less than 68%) compared with the other subfamily members.

A combination of *in vitro* and *in vivo* studies showed that UGT73C6 can glycosylate flavonoids (Jones et al., 2003). In addition, UGT73C5 and UGT73C6 were indicated to catalyze the *in vivo* 23-*O*-glycosylation of brassinosteroids (Poppenberger et al., 2005; Husar et al., 2011), *i.e*., plant steroid hormones that promote growth by modulating cell division, elongation, and differentiation. Glycosylation leads to inactivation of the hormone and thus, is important, together with hydroxylation, to maintain cellular homeostasis. Overexpression of *UGT73C6* or *UGT73C5* in *A. thaliana* accordingly results in phenotypes indicative of brassinosteroid deficiency, with plants showing a cabbage-like morphology and dark green leaves with short petioles. In addition, and in correlation with overexpression strength, delayed flowering and senescence, and a reduction in fertility were observed (Poppenberger et al., 2005; Husar et al., 2011).

Leaf size and shape are key determinants of plantś ability to absorb sunlight and convert it into chemical energy through photosynthesis (Gonzalez et al., 2012). Leaf growth is a dynamic process regulated by two key, highly interrelated processes, namely cell division and cell expansion (Kalve et al., 2014). In Arabidopsis, founder cells located at the sides of the shoot apical meristem initiate leaf development (Vercruysse et al., 2020). After the initial phase of extensive cell proliferation throughout the leaf primordium, cell division ceases and further growth is supported mainly by cell expansion (Gonzalez et al., 2012). The latter process initiates at the leaf tip and follows a basipetal direction (Andriankaja et al., 2012). Accurate regulation of the cell proliferation and cell expansion mechanisms is therefore critical for determining final leaf size. Six gene modules are involved in the regulation of Arabidopsis leaf growth and one of them, the *GROWTH REGULATORY FACTOR* (*GRF*)-*GRF-INTERACTING FACTOR* (*GIF*) module, plays an important role in determining the cell number in leaves by controlling cell proliferation (Vercruysse et al., 2020).

In the present work, we show that alterations in the transcript levels of two *UGT73C6*- overlapping lncNATs (*NATs_UGT73C6_*) lead to changes in the size of the *A. thaliana* rosette. Microscopical analyses indicate that these differences are mainly due to modifications in the number of cells suggesting that the levels of *NATs_UGT73C6_* affect cell proliferation. These phenotypic effects were demonstrated to be due to *NATs_UGT73C6_* acting as *bona fide* noncoding RNAs and were not caused by the activity of small, RNA-encoded peptides. Our data suggest that the *NATs_UGT73C6_* exert their function in *trans* and independently of the sense gene in a mechanism that excludes downregulation of the *UGT73C* subfamily members but involves the regulation of important transcription factors such as *GRFs* and *GIFs*.

## Results

*NAT1_UGT73C6_* and *NAT2_UGT73C6_*, two lncRNAs overlapping the *UGT73C6* gene, are expressed in diverse organs and at different developmental stages of *A. thaliana* Analysing the transcripts reported in the TAIR10 database, we identified two natural antisense transcripts (*At2g36792.1* and *At2g36792.2*) that overlap the *UGT73C6* gene. *In silico* analysis of the protein-coding capacity of both transcripts, hereafter referred to as *NAT1_UGT73C6_* and *NAT2_UGT73C6_* (for *At2g36792.1* and *At2g36792.2*, respectively) and collectively *NATs_UGT73C6_*, revealed the presence of several open reading frames (ORFs) potentially encoding peptides of at least five amino acids. Most of the predicted peptides were smaller than 30 amino acids in length and none of them showed similarity to peptides present in the databases, nor did they have functional domains as determined using Pfam and PROSITE. Together, these data suggest that the transcripts are *bona fide* lncNATs.

To analyze the expression pattern of the individual *NATs_UGT73C6_*, we generated transgenic lines in which the *uidA* gene (*GUS*), encoding β-glucoronidase (GUS) was placed under the control of 2054 and 2580 bp sequences upstream of the annotated transcription starts of *NAT1_UGT73C6_* and *NAT2_UGT73C6_*, respectively, which were expected to contain the respective NATs promoters (Figure 1A). Analysis of *GUS* expression in several independent homozygous lines (three for *NAT1_UGT73C6_:GUS* and five for *NAT2_UGT73C6_:GUS*) revealed that both promoters were active and, interestingly, these activities were independent of the presence of the sense gene at the same locus. The selected promoters showed a distinct pattern of expression: the *NAT1_UGT73C6_* promoter was active exclusively in roots, whereas the activity of the *NAT2_UGT73C6_* promoter was detected only in the aerial organs, particularly in young leaves (Figure 1B). As the selected sequences overlap completely except for 526 bp at the 3’ end (Figure 1A) these results suggest that motifs present in the latter region are responsible for the expression of the *NAT2_UGT73C6_* promoter in aerial tissues. Subsequent time course experiments confirmed that the promoter activity of both antisense transcripts is restricted to the aforementioned organs at all the analyzed developmental stages (Supplemental Figure 1A).

**Figure 1.**
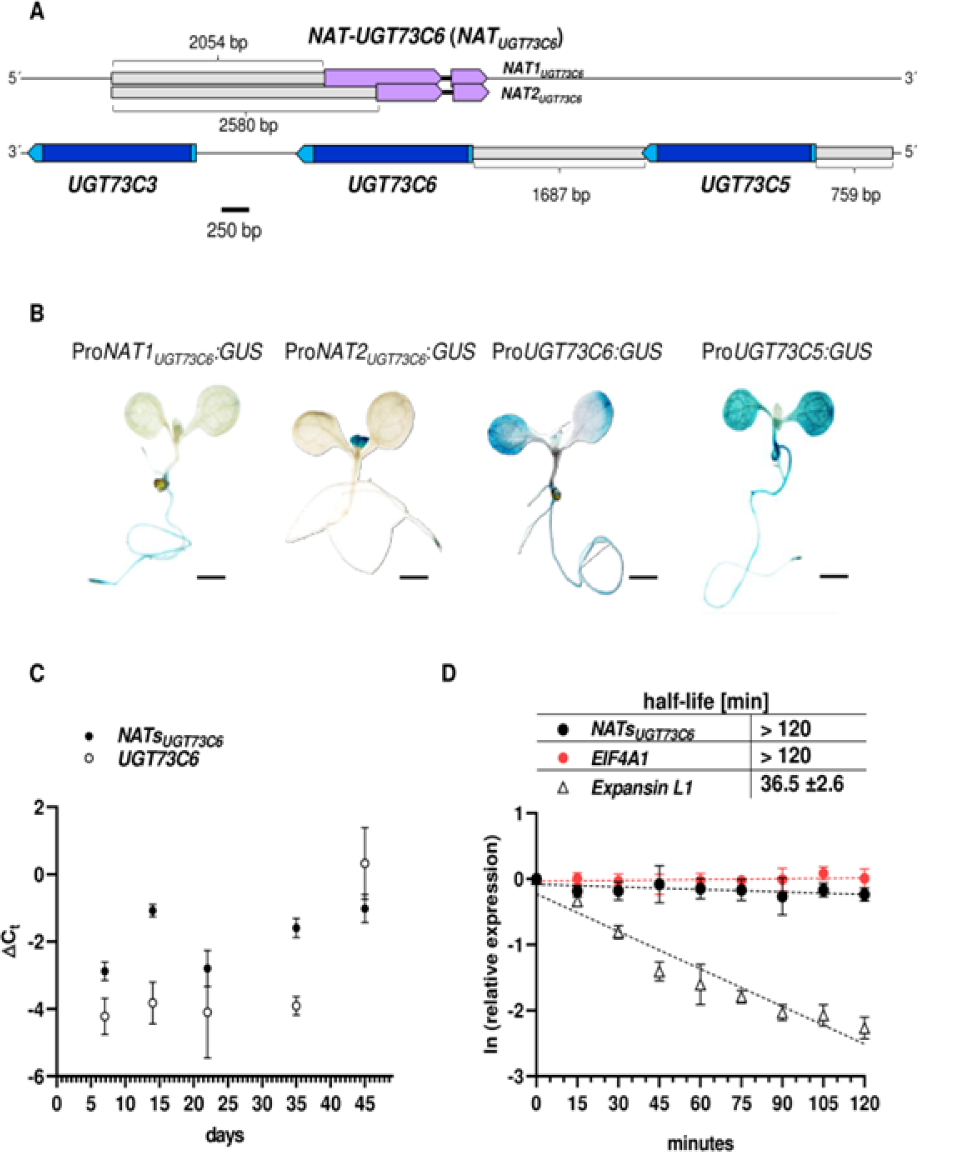
NATs_*UGT73C6*_ are stable and expressed at different developmental stages. (A) Schematic showing the geoomic arrangement of *UGT73C6* and its antisense gene NAT_*UGT73C6*_ and the llanking protein-cooing genes *UGT73C3* and *UGT73C5*. The regions used as promotocs 10 direct the expression of the GUS *(β-glucuronidase)* reporter gene are represented by grey boxes and their lengths indicated in base pairs (bp). Protein-coding genes *(UGT73C6, UGT73C5and UGT73C3)* and the natural antisense long non-coding ANAs cooing gene *NAT0017JC6 are* indicated with blue and violet boxes, respectively. Untranslated regions (UTAs) are represented by light blue boxes and intronsby thin black lines. The region, on the chromosome 2 of *A. thaliana*, is represenled in scale. (B) Histochemical analysis ot GUS activity in 7-day-old seedlings grown on 1)4ates under long­ day (16 h lighV 8 h dark cycle) conditions. Representative homozygous lines were chosen for the detection ol the promoter activity of NAT1_*UGT73C6*_ (ProNAT1_*UGT73C6*_GUS), UGT73C6 (ProUGT73C6:GUS), and *UGT73C5* (Pro*UGT73C5.GUS*),. Seedlings were stained overnight. Bars = 5 mm. (C) Reverse transcription quantita1ive PCA (RT-qPCA) expression profiling of NATs_*UGT73C6*_, and *UGT73C6* transcripls at various developmental stages. Seedlings and plants were grown under long-day conditions. AT-qPCA was pertormed on three to live independent biological replicates (pools of seedlings grown on plates for 7 and 14 days, and ol 4 10 10 rosettes withou1 floral stems of plants gro’Nf’I on soil for 22, 35 and 45 days}. Data are means with SD from ΔC_t_, values (C_t_, reference gene *PP2A* - C, gene of interest). (D) NATs_*UGT73C6*_, RNA stability in wild-type (Col-OJ seedlings. The half-life (t_1/2_, Table) *was* calculaled based on degradation curves a er cordycepin trealment (plot). Stable *(Eukaryotic translation initiation factor 4AI, EIF4At)* and short-lived *(Expansin like* 1, *Expansin L1*) transcripts were inciuded as controls. Values correspond to means ol three independent experiments wilh error bars representing SD. NATs_*UGT73C6*_ include both, NAT1_*UGT73C6*_ and NAT2_*UGT73C6*_, transcripts.

To compare the expression domains of the sense and antisense genes we next generated and tested reporter lines for *UGT73C6* and, additionally, for its closest w7 e observed *UGT73C6* promoter activity in roots and cotyledons (Figure 1B) but not in young leaves where *NAT2_UGT73C6_* is detected, suggesting a spatiotemporal separation of the expression of the sense and antisense genes in these organs. Analyses of the *UGT73C5* promoter activity revealed a strong expression in cotyledons and roots, where its expression overlaps with those of *NAT1_UGT73C6_* and *UGT73C6*, but not in young developing leaves (Figure 1B). The *NAT1_UGT73C6_* promoter was not active in flowers whereas *NAT2_UGT73C6_* promoter activity was detected in stamen filaments and in silique pedicels (Supplemental Figure 1B). The promoters of *UGT73C6* and *UGT73C5* were found to be active in anthers and silique pedicels, and in sepals, stamen filaments and style apex, respectively (Supplemental Figure 1B).

Using RT-qPCR we also determined the expression of the sense and antisense genes at different time points (Figure 1C). The data suggest that the *NATs_UGT73C6_* (mainly *NAT2_UGT73C6_* according to the pattern detected with the reporter gene line showing activity in aerial organs) were expressed at all analyzed times. Except for the last time point included in the study (45 days after stratification, DAS), the transcript levels of *NATs_UGT73C6_* were observed to be higher than those of *UGT73C6* (Figure 1C).

Altogether, these results suggest that the promoters of both *NATs_UGT73C6_* transcripts are active, that the expression of the RNAs is developmentally regulated and that it occurs independently of the transcriptional activity of the *UGT73C6* gene in *cis*.

### *NATs_UGT73C6_* transcripts are alternatively spliced and stable

According to the TAIR10 database the transcripts encoded by the *NAT_UGT73C6_* gene contain an intron (Figure 1A). To determine the splicing pattern of the *NATs_UGT73C6_* transcripts we extracted total RNA from shoots and roots of 9-day-old seedlings grown in long day conditions and performed RT-PCR. Whereas *NAT1_UGT73C6_* was hardly detected over cDNA from roots, a clear amplification of *NAT2_UGT73C6_* was observed over cDNA in shoots (Supplemental Figure 2A). The NAT2_UGT73C6_ amplicon was cloned and analysis of individual clones by colony PCR revealed the presence of inserts of different sizes (Supplemental Figure 2B). Sequence analyses of the clones demonstrated that the *NAT2_UGT73C6_* transcripts consist of splicing variants in which half of the population retains the intron. The other half of the transcripts was shown to consist of three splicing variants with lengths of 1013, 1005 and 985 nucleotides, which are suggested to be generated using alternative, canonical splicing donor and acceptor sites (Supplemental Figure 2C).

The presence of fully spliced variants indicates that at least a fraction of the *NATs_UGT73C6_* transcripts is transported to the cytosol. Considering that nuclearly localized lncRNAs are often unstable (Clark et al., 2012) whereas lncRNAs with a *trans*-acting mode of action in the cytosol are more stable, we decided to determine the half-life of the *NATs_UGT73C6_* transcripts. With this aim in mind, we treated seedlings with cordycepin, an adenosine analogue that induces termination of chain elongation when incorporated during RNA synthesis and quantified the transcript abundance at different time points, as previously described (Fedak et al., 2016). Stability determination based on degradation curves showed that the *NATs_UGT73C6_* transcripts have a relatively long half-life (>120 min). Actually, the half-life of the *NATs_UGT73C6_* transcripts was found to be comparable to that of the stable protein-coding transcript of a housekeeping gene (*Eukaryotic translation initiation factor 4A1*, *EIF4A*) and longer than that of a short-lived mRNA (*EXPANSIN-LIKE1*, *Expansin L1*) (Figure 1D). In addition, preliminary data from subcellular fractionation analyses suggested that the *NATs_UGT73C6_* mainly localize to the cytosol (data not shown).

Overall, these data showed that the *NATs_UGT73C6_* transcripts are alternatively spliced and exhibit relatively high stability suggesting that they are transported to the cytosol.

### *NATs_UGT73C6_* downregulation leads to a reduction in the rosette area by decreasing leaf size and the number of mesophyll and epidermal cells

Considering a potential *trans*-activity of the *NATs_UGT73C6_* transcripts, we decided to analyze the effects of their downregulation. To achieve this, we first tested the earlier reported *ugt73c6_ko_* mutant line (SAIL_525_H07) (Jones et al., 2003), which contains a T-DNA insertion at position 1440 downstream of the reported *UGT73C6* transcription initiation site (Supplemental Figure 3A). The insertion generates a premature stop codon at position 472 leading to a protein that is truncated C-terminally by 24 residues. RT-qPCR analysis confirmed that, *UGT73C6* expression is strongly reduced in the mutant line. However, unexpectedly, the *NATs_UGT73C6_* levels turned out to be unaltered (Supplemental Figure 3B), suggesting that not only the transcription of *NAT2_UGT73C6_* but also that of *NAT1_UGT73C6_* can be initiated downstream of the T-DNA insertion. This line thus turned out to be unsuitable to study the effects of *NATs_UGT73C6_* downregulation.

We therefore generated two different collections of transgenic lines, which, *via* the *35S* promoter overexpress artificial microRNAs (amiRNAs) (Carbonell et al., 2014) that specifically target both *NAT_UGT73C6_* transcripts (Supplemental Figure 4A). In the first set of 15 independent homozygous lines, three were confirmed unambiguously to express the amiRNAs (Supplemental Figure 4B). Interestingly, these plants showed a decrease in rosette size when compared to wild-type plants and to plants transformed with the empty vector. The *NATs_UGT73C6_* levels of these lines were found to show a reduction close to or higher than 50 percent (lines 1, 2 and 5, Supplemental Figure 4C) as compared to the control plants when analyzed by RT-qPCR.

To quantify the observed phenotypic differences we analyzed 25-day-old plants using an available software (Easlon and Bloom, 2014). This confirmed the initial, macroscopic impression that lines indeed showed a reduced rosette area in comparison to the control plants (Supplemental Figure 4E). With the second set of lines, which was generated afterwards, we first analyzed the levels of *NATs_UGT73C6_* in eight independent homozygous lines and identified one (*_amiR_NATs_UGT73C6_-10*) in which these were strongly reduced (Supplemental Figure 4D). Interestingly, and consistent with the previous results, the plants from this line were the only ones that showed smaller rosettes when compared with wild-type plants and plants overexpressing amiRNAs targeting the *GUS* gene used as controls (Figures 2A and B). This phenotype was confirmed by further experiments (Figure 2C). Compiled data from each of these independent experiments showed that the average decrease in rosette area in lines with reduced *NATs_UGT73C6_* levels range between 13 to 20 percent (Figure 2C). In accordance with this, plants of the *_amiR_NATs_UGT73C6_-7* line and of the other lines (*_amiR_NATs_UGT73C6_-8* to *-9* and *-11* to *-14*), in which the *NATs_UGT73C6_* levels were not altered showed no differences in rosette size (Figure 2C and Supplemental Figure 4F). In sum, we took these data as an indication that reduction in *NATs_UGT73C6_* transcripts leads to a decrease in the rosette area.

**Figure 2.**
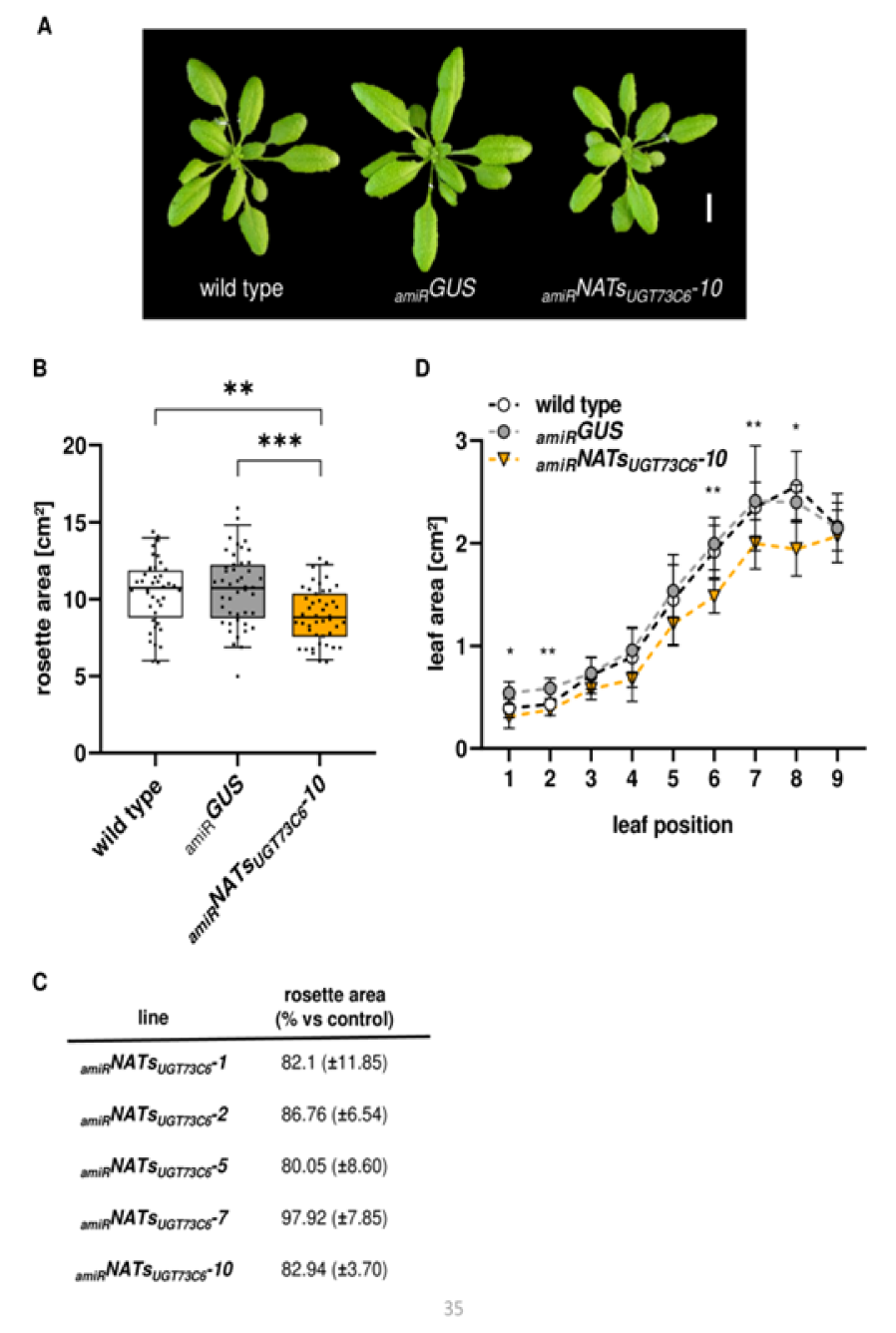
Effects of NATs_*UGT73C6*_, downregulation on roselle area and leaf size. (A) Representative pictures or 25-day-old wild-type) (left), _*amR*_GUS control (center) and _*amR*_NATs_*UGT73C6*_ 10(right) plants. Bar = 1 cm. (B) Rosette area of _*amR*_NATs_*UGT73C6*_ plants compared to wild-type and control plants expressing a GUS-specific amiRNA(_amR_GUS). Plants were grown under long-day (16 h light/ 8 h dark cycle) conditions and photographed at 25 days alter sowing (DAS). Shown are compiled data from two Independent experiments including 20 to 25 plants per experiment. Box plots show medians and interquartile ranges, whiskers extend from 5 to 95 percentile and values from individual rosettes are shown as black dots. Asterisks denote statistical differences from the wild type at p ≤ O.001 (**) and p ≤ 0.001 (***) assessed by one-way ANOVA and Tukey’s multiple comparison test. (C) Ouanlification of rosette area reduction in plants from _*amR*_NATs_*UGT73C6*_ expressing lines compared to control plants (translonned with the empty vec1or or expressing _amR_GuS). Plants from the _*amR*_NATs_*UGT73C6*_ line, which shows wild-type levels of NATs_*UGT73C6*_, were included as an additional control. Numbers are means with SD of means from lndividual experiments expressed as percentage compared to the control (100%). Data correspond to two _*amR*_NATs_*UGT73C6*_, three (_*amR*_NATs_*UGT73C6*_-1, _*amR*_NATs_*UGT73C6*_-5) and four (_*amR*_NATs_*UGT73C6*_-10) independent experiments. Including 18 to 36 plants each. (D) Area of individual leaves of 30-day-old wild-type, _*amR*_GUS and _*amR*_NATs_*UGT73C6*_-10 plants grown in soil. The graph shows mean and SD (n =4.5) of areas from leaves 1 to 9. Aslerisks Indicate statistical differences from the wild type at p ≤ 0.05(*) and p ≤ 0.01 (**) assessed by IWO· tailed S1udenl’s *t*-Test.

To exclude a potential effect of the UGT73C6 protein on the plant phenotype, we also performed rosette area measurements in the earlier mentioned *ugt73c6_ko_* line. The obtained data demonstrated that the absence of a functional UGT73C6 protein has no influence on the rosette size (Supplemental Figure 3C), reinforcing the idea that the observed phenotypic changes result from the reduction of *NATs_UGT73C6_* levels.

To understand the contribution of the individual leaves to the overall change in rosette size we determined the area of individual leaves from 30-day-old plants. Our results showed that all the leaves from the *_amiR_NAT*1*s*1*_UGT73C6_-10* plants were smaller than those of the control plants, with differences that were statistically significant for leaves 1, 2, 6, 7 and 8 (Figure 2D). In a second experiment, we quantified the growth of individual leaves over time. We observed that the leaves from plants of the *_amiR_NATs_UGT73C6_-10* line were smaller at all time points analyzed (Supplemental Figure 4G) and that the differences were maintained even after leaf growth was completed (*e.g.*, leaves 1, 2, 3 and 4, Supplemental Figure 4G). These results suggested that a reduction in size of the individual leaves was responsible for the smaller rosette area in the lines with reduced *NATs_UGT73C6_* levels.

We next wanted to understand whether the observed phenotype of the lines with downregulated *NATs_UGT73C6_* levels was possibly caused by changes in cell proliferation or in cell expansion. For this purpose, we determined the cell size of palisade mesophyll and abaxial epidermal cells from the tip and the bottom regions of the leaf 6 from *_amiR_NATs_UGT73C6_-10* and from the control (*_amiR_GUS*) plants. Combined data including the cells from both leaf regions showed no differences in mesophyll cell size but a statistically significant increase in epidermal cell area in the *_amiR_NATs_UGT73C6_-10* line (Figure 3B). Using these data, we estimated the total number of cells by dividing the determined total area of the individual leaves (Figure 3A) by the average cell area of each cellular type (Figure 3C). Results of these estimations showed that the total number of both cell types were significantly lower in the lines where the *NATs_UGT73C6_* levels had been reduced by the amiRNA knockdown approach (Figures 3C and 3D).

**Figure 3.**
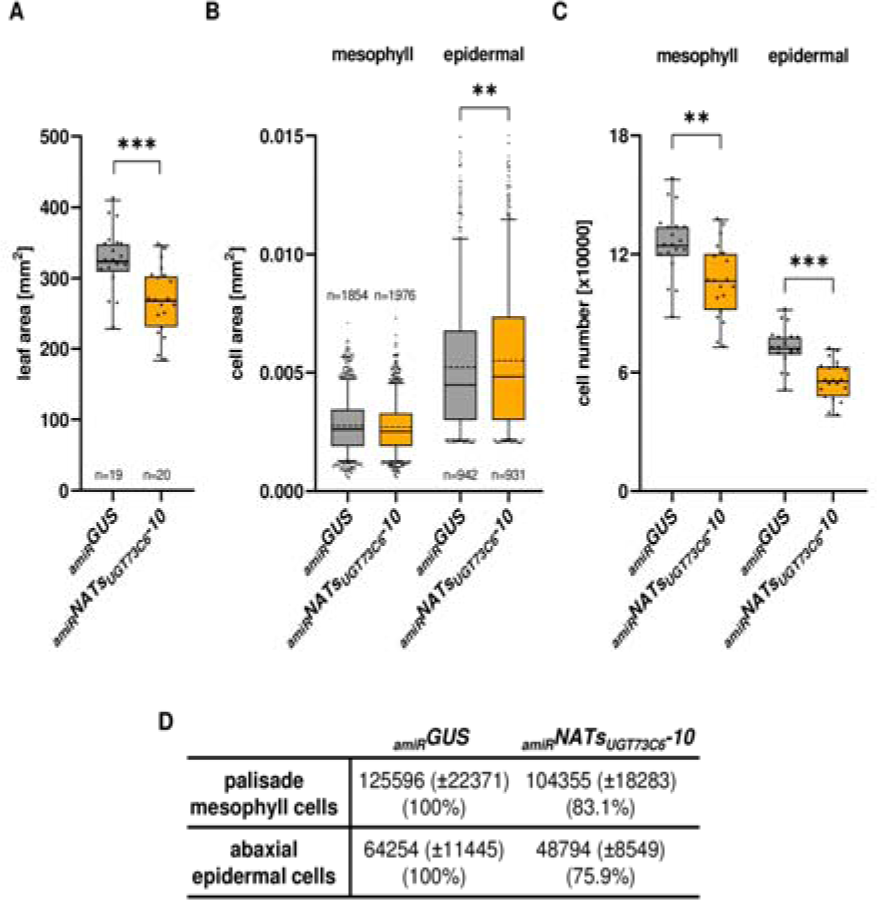
NATs_*UGT73C6*_ downregulation has a negative effect on cell number. (A) Leaf area, (B) mesophyi and epidermal cell area, and (C) estimated mesophyill and epidermal cell number of the leaf 6 from 35-day-old _*amR*_NATs_*UGT73C6*_-10 and control (_amR_GUS) plants. Estimated lolal cell number was obtained dividing areas from lnddividals (n = 19.20) by the average cell area. Box plots show medians (continuous line) and inte,quanile ranges, whiskers extend from 5 to 95 percentile. Dots represeof lndividua values (A and C). ln (B) dotted lines and tilled dots indicale average cell area and outliers. respectively. Unpaired, two-tailed Student’s *t*-test applied for leaf area and cell number eslimation, and one-way ANOVA followed by Tukey’s multiple comparsions test for cell area. Asterisks. indicate statistical difference from the control at p < 0.05 (*), p < O.01 (**) and p < 0.001 (***). n indicates the number of measured leaves (A) or cells (B). (D) Estimated total cell number in leaf 6 of 35-day-old _*amR*_NATs_*UGT73C6*_-10 and control (_*amR*_GUS) plants. Values are means with SD of means from individual leaves (n = 19-20) and percentages compared to the control (between parentheses). Abaxial epidermal cells bigger than 0.015 mm^2^, composing less than four percent of the measured cells., were excluded from the analysis.

These data suggest that the smaller size of leaf 6 of the *_amiR_NATs_UGT73C6_-10* plants is mainly due to a decrease in cell number. Considering that in this line all leaves were found to be smaller than those of the control plants, we explained the reduction in rosette area by a reduction of the number of cells in each leaf. This idea was supported by the fact that in the plants with regulated *NATs_UGT73C6_* levels the estimated reduction of cell number (ca. 20% if both cell types are considered) correlated closely with the observed reduction in rosette area (17%) (Figures 3D and 2C, respectively).

### Overexpression of *NAT1_UGT73C6_* or *NAT2_UGT73C6_* induces an increase in rosette size

Considering that the previously described phenotype was a consequence of *NATs_UGT73C6_* downregulation, we now performed an experiment to determine if an increase of the *NATs_UGT73C6_* levels would lead to the opposite phenotype. For this purpose, we generated transgenic lines expressing each of the *NATs_UGT73C6_* under the control of the *35S* promoter. Several independent homoz1y3gous lines were selected (4 overexpressing *NAT1_UGT73C6_* and 4 overexpressing *NAT2_UG_*1*_T_*4*_73C6_*, Supplemental Figures 5A and 5B) and rosette area determination was performed as described before. In all cases, plants from transgenic lines overexpressing the *NATs_UGT73C6_* sequences showed a small but significant increase in rosette size compared to control plants (Figure 4). Analyses from at least three independent experiments confirmed that overexpression of either *NAT1_UGT73C6_* or *NAT2_UGT73C6_* leads to an increase in rosette size, suggesting that, indeed, high levels of *NATs_UGT73C6_* transcripts determine this phenotype (Figure 4C). While lines overexpressing *NAT1_UGT73C6_* showed an increase of the rosette size ranging between 10 and 23 percent, these values were slightly higher in lines overexpressing *NAT2_UGT73C6_* (14 to 32 percent) (Figure 4C). If all lines were considered the average rosette increase was 14.65 (±5.52) percent for *NAT1_UGT73C6_* and 22 (±7.47) percent for *NAT2_UGT73C6_*. Plants overexpressing *UGT73C5* included as a control showed the reported brassinosteroid-deficiency phenotypes and a strong reduction (more than 40 percent) of the rosette area (Figures 4A and 4C).

**Figure 4.**
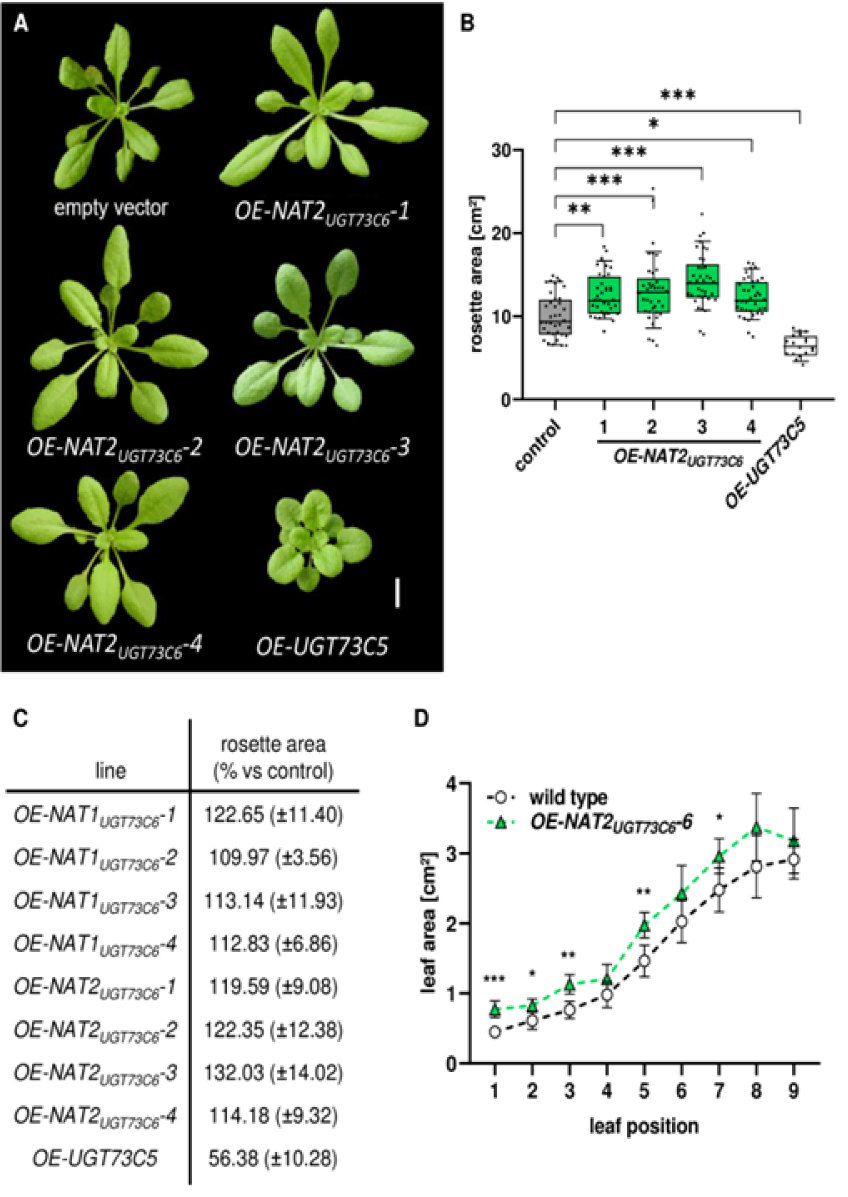
NATs_*UGT73C6*_ overexp ression increases rosette area. (A) Pictures from 25-day-old control (transformed with the empty vector) and NAT2_*UGT73C6*_ overexpressing (OE) plants. Representative plants corresponding to four independent homozygous lines are shown. Plants overexpressing *UGT73CS* were inciuded for comparison. Bar = 1 cm. (B) Roselle area of NAT2_*UGT73C6*_ Overexpressing (OE) plants compared to control plants transformed with the empty veclor. Plants were grown on soil under long-day (16 h light/ 8 h dark cycle) condtions and photographed at 25 DAS. Compiled data of two independent experiments including 18 and 19 planls, respectively. Planls overexpressing *UGT73C5* (n = 18) were Included as controls in one experiment. Box plots show medians and interquartile ranges, whiskers extend from 5 to 95 percentile and individual values are shown as black dots. Asterisks denote statistical significance at p ≤ 0.05 (*). p ≤ 0.01 (**) and p ≤ 0.001 (***) assessed by one-way IWOVA and Tukey’s multiple comparison lest. (C) Quantilica1ion of the rosette area in homozygous lines overexpcessing NAT1_*UGT73C6*_ or NAT2_*UGT73C6*_ compared to a control line transformed with the empty vector. Numbers are means with SD of means from individual experiments expressed as percentage of the control (set at 100%). Data correspond to three independent expenments including 17 to 20 plants each. (D) Area of individual leaves of 30-day-old wild-type and NAT2_*UGT73C6*_ overexpressing (OE-NAT2_*UGT73C6*_-6) plants grown in soil. The graph shows mean and SD (n = 4.5) of areas from leaves 1 to 9. Asterisks indicale statistical differences from the wild type at p ≤ 0.05 (*), p ≤ 0.01 (**) and p<0.001 (***) assessed by two-tailed Student’s t-Test.

**Figure 5.**
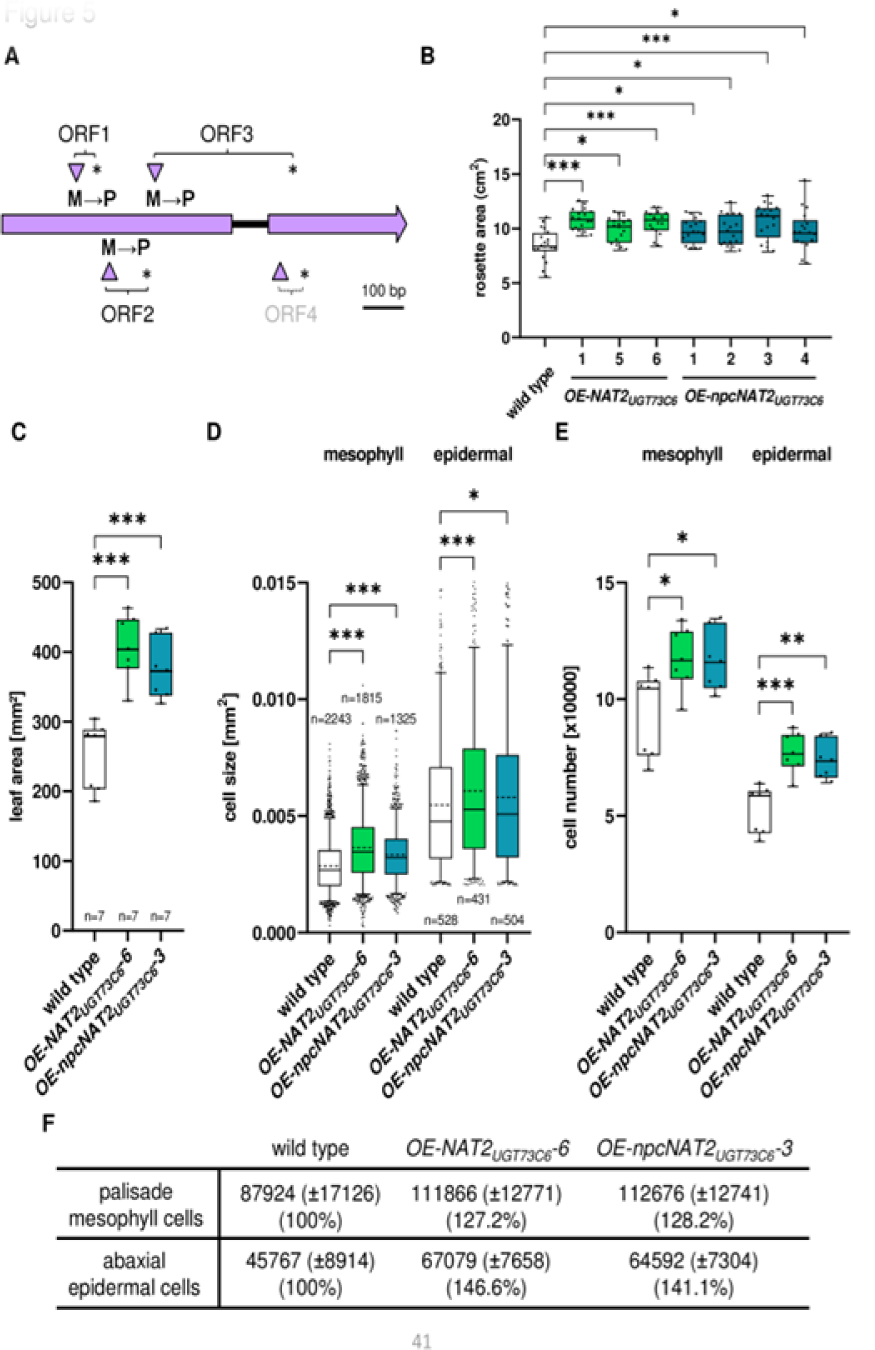
Overexpression of NAT2_*UGT73C6*_ or of the non-peptide coding NAT2_*UGT73C6*_, variant increases rosette and leaf area and the number of epidermal and mesophyil cells. (A) Schematic of the non-peptide coding NAT2_*UGT73C6*_(npcNAT2_*UGT73C6*_) variant. Methionine to proline (M→P) changes introduced by site-directed mutagenesis of the start codons at positions 198 (ORF1). 289 (ORF2) and 405 (ORF3) are indicaled. Arrowheads and asterisks indicate start and slop codons, respectively. (B) Roselle area of wild-type plants and plants overexpressing NAT2_*UGT73C6*_(OE-NAT2_*UGT73C6*_) and the non-peptide coding NAT2_*UGT73C6*_, variant (OE-NAT2_*UGT73C6*_) Plants were grown on soil under long-day (16h light/ 8 h dark cycle) conditions and photographed at 25 DAS. Shown are data from one representative experiment lnclucding 19 and 17 plants for wild-type and transgenic lines, respectively. Box plots show medians and interquartile ranges; whiskers extend from 5 to 95 percentile. Asterisks denote sinificant statistical differences at p ≤ 0.05 (*), p ≤ 0.01 and p ≤ 0.001 (***) assessed by one-way ANOVA and Tukey’s multiple comparison lest. (C) Leaf area. (D) mesophyll and epidermal cell area, and (E) estimated mesophyll and epidermal cell number of sixth leaves from 35-day-old control plants (wild type) and plants overexpressing NAT2_*UGT73C6*_, (OE-NAT2_*UGT73C6*_-6) or the non-peptide coding NAT2_*UGT73C6*_, variant *(OE-NAT2*_*UGT73C6*_-3 Estimated total cell number was obltained dividing the area of the individual leaves by the average cell area for each cell type. Box plots show medians (continuous line) and Interquartile ranges, whiskers extend from 5 to 95 percentile. Dots represent individual values in (C) and (E). Doned lines and filled dots in (D) indicate average cell area and outliers, respectively. Asterisks denote significant slatistical differences at p ≤ 0.05 (‚), p ≤ 0.01 (**) and p ≤ 0.001 (***) assessed by unpaired, two-tailed S1udenrs nest for (C) and (E), and by one-way AfiOVA and Tukey-s multiplecomparison test (D). n Indicates the number of measured leaves (C) or cells (D). (F) Estimated tolal cell number in leaf 6 of 35-day-old wild type, OE-NAT2_*UGT73C6*_-6 and OE-NAT2_*UGT73C6*_-3 plants. Numbers are means with SD of means from individual leaves (n = 7). Percentages versus control (100%) are indicated between brackets. Abaxial epidermal cells larger than O.015 mm^2^, including less than six percent of the measured cells, were removed from the analysis.

To determine again the contribution of individual leaves to the changes in rosette area, we dissected and measured leaves from 30-day-old *NAT2_UGT73C6_*-overexpressing plants and wild-type plants. Our analysis revealed that leaves of plants overexpressing *NAT2_UGT73C6_* (*OE-NAT2_UGT73C6_-6*, a new generated line with high *NAT2_UGT73C6_* expression levels, Supplemental Figure 8) were indeed larger than those of the control plants, showing statistically significant differences for leaves 1, 2, 3, 5 and 7 (Figure 4D). In addition, we quantified individual leaf growth by measuring leaf size at different time points. The analysis of the growth curve showed that the area of the individual leaves of the overexpression line was larger than that of the corresponding leaves of the control plants (Supplemental Figure 5C). Furthermore, we observed that the size differences also remained when leaf development was completed, suggesting that these differences did not result from a faster growth of the *NAT2_UGT73C6_* overexpression line.

Altogether, these data support the idea that the overexpression of *NATs_UGT73C6_* induces an increase in the size of the individual leaves and thus an enlargement of the rosette area.

### Increase in rosette area is caused by a *bona fide* lncRNA activity of the *NATsUGT73C6*

According to the current definition, lncRNAs should not contain ORFs encoding for peptides of more than 70 amino acids (Ben Amor et al., 2009). As already explained above, most of the ORFs predicted in the *NATs_UGT73C6_* encode less than 30 amino acids. However, *NAT1_UGT73C6_* and *NAT2_UGT73C6_* encode for a 103 amino acids long polypeptide, peptide 3 (Supplemental Figure 6A). Thus, it was important to determine whether these peptides are produced *in planta* and to understand whether the earlier phenotypes observed in the knockdown and overexpression lines were possibly associated with activities of these peptides. As overexpression of either *NATs_UGT73C6_* causes a similar phenotype in the transgenic plants, we decided to perform the following experiments solely with *NAT2_UGT73C6_*.

First, to enable and visualize expression of the four longest ORFs that can be hypothesized in the sequence of both *NATs_UGT73C6_* (Supplemental Figure 6A), we generated C-terminal GFP-fusion constructs by removing the translational stop codons from the ORFs by site-directed mutagenesis. In one set of constructs, we kept the *NAT2_UGT73C6_* sequence upstream of the individual ORFs to preserve potential regulatory translation initiation elements in the transcripts (Supplemental Figure 6B, right). In a second set of constructs, expression of the ORFs was enabled only via individual start codons (Supplemental Figure 6B, left). Transient expression assays, which were performed in *N. benthamiana* and in the presence of the viral silencing suppressor p19 (Scholthof, 2006) to minimize effects of posttranscriptional gene silencing, revealed expression of most of the fusion peptides with both sets of constructs (Supplemental Figures 6C and 6D). An exception was the fusion peptide corresponding to ORF4. This was only detected in leaves that were infiltrated with the construct allowing the expression of the individual ORF (Supplemental Figures 6C and 6D). Based on this finding we focused the further analysis on ORFs 1 to 3, which anyhow cover most of the peptide-coding *NAT2_UGT73C6_* sequence.

**Figure 6.**
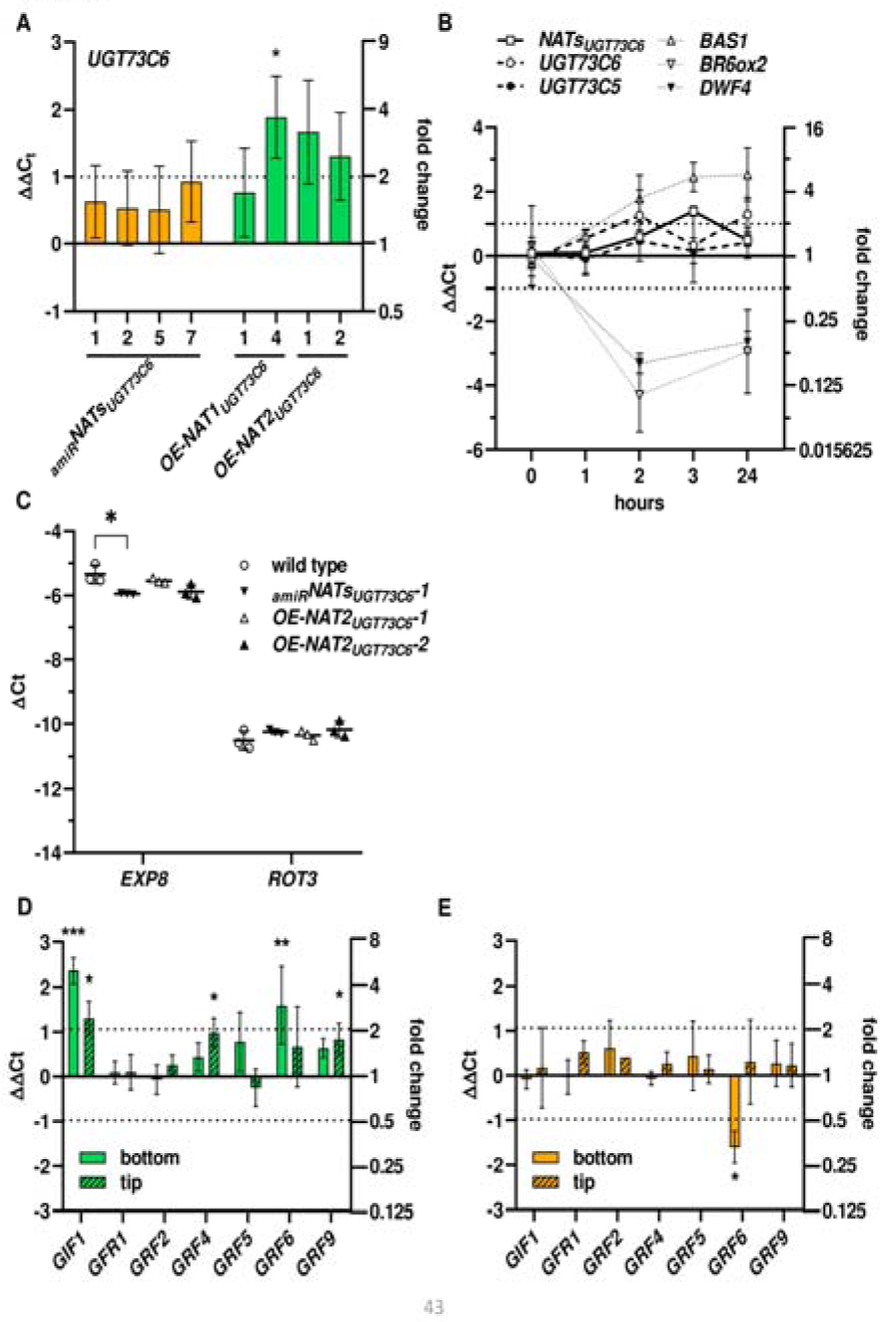
Levels of *GRFs* but not of *UGT73C6* or brassinosteroid marker genes are affected In lines with altered NATs_*UGT73C6*_, expresstion. (A) *UGT73C6* levels determined by RT-qPCR in lines with reduced (_amR_) and enhanced (OE) NATs_*UGT73C6*_ expression. Seedlings were grown under long-day (16 h light/ 8 h dark Cycle) conditions for 14 days. Bar plots show ΔΔC_t_ [mean difference in ΔC_t_ between lines with modified incNATs expression and control line (ΔC_t_ amiRNA expressing line or NATs_*UGT73C6*_ overexpressing line - ΔC_t_ control line) and staooard error of mean differences of three lndependent bilogical replicates (pools of seedings)]. Expresion levels normalized to the reference gene *PP2A* Asterisks denete sigmlicant stastical difference at p < 0.05 (*) assessed by unpaired, two-tailed Student’s *t*-Test. (B) Response of NATs_*UGT73C6*_ and *UGT73C5* expression to brassrnosteroid tratment determined by RT-qPCR. Seven-days-old seedlings grown in liquid media were treated with either 1μm 24-epiBL or DMSO (moc, treatment) and samples were collected at the indicated time points. Compiled data from independent experiments including at least three biological replicates (pools ol seedlings). Symbols lndicate (ΔC_t_, 24-epiBl-treated - ΓC_t_ DMSO-treated samples)] and standard error of the mean differences. Expression analysis of the Bl-responsive genes *PHYROCHROME P450 9081 (DWF4), BRASStNOSTERD-6-0XIDASE 2* (*BR6o’2*) and *PHYB ACnVATION- TAGGED SUPRESSOR 1* (*BASI*) was included to monitor treatment effectiveness. Expression levels were normalized to the reference gene *GAPC2* Dotted lines indicate two-fold changes (C) Expression of the B-R respansive genes *ROTUNDIFOLIA3 (ROT3)* and *EXPANSIN 8 (EXP8).* in transgenic lines with reduced (_amR_) or enhanced (OE)NAT_*UGT73C6*_ expression. Depicted are means (tick lines) with error bars showing SD of ΔC_t_, values (C_t_, reference gene *GAPC2* - C, gene of imerest) of three bioiogical replcates (independent pools of 10-day-old seedlings grown vertically on planes under long-day conditions). Asterisks indicale significant Stastical differenceat p ≤ 0.05 (*) assessed by unpaired, two-tailed Students *t*-Test. (D) and (E) *GIf1* and *GRFs* expression analysis at tip and bottom regions of third leaves from lines with enhanoced (OE-NAT2_*UGT73C6*_-6) or reduced (_amR_NATs_*UGT73C6*_-10) NATs_*UGT73C6*_ expression compared with control lines. Bar plots show ΔΔC_t_, values (mean difference in ΔC_t_ between lines with altred NATs_*UGT73C6*_ expression and control (ΔC_t_, in NAT2_*UGT73C6*_-6, or _amR_NATs_*UGT73C6*_-10 ΓC_t_ in control)] and standard error of mean diffrence of three to four biologlcal replicates (independent pools of leaf sections). Expression levels were normalized the reference gene *PP2A*. Plain and striped bars indicate values of bottom and tip regions, respectively. Dotted lines indicate two-fold changes. Asterisks denote significant statistical differences at p ≤ 0.05 (*). p ≤ 0.01 (**) and;, ≤ 0.001 (***) assessed by unpaired. two-taied Student’s *t*-Test. Leaves were collected from 12-day-old seedlings grown on plates under long-day conditions.

To understand whether the phenotype of increased rosette area derives from the action of the *NAT2_UGT73C6_*-encoded peptides, we generated transgenic *A. thaliana* lines, which comprised the earlier tested constructs allowing the expression of either ORF3 alone or of ORFs1 to 3 including the corresponding *NAT2_UGT73C6_* upstream sequences (Supplemental Figures 7A and 7B). The expression levels of the selected homozygous lines were quantified (Supplemental Figures 7A and 7B) and the rosette area of plants measured. As shown in Supplemental Figure 7C, none of these transgenic lines showed a significant increase in rosette area. Moreover, some lines actually grew to even smaller sizes than the control plants.

These data suggested that the phenotypes that were observed in the earlier experiments were not caused by the activity of the peptides encoded by *NAT2_UGT73C6_*.

A second set of transgenic lines was generated to determine whether the phenotypes that we observed during the overexpression of *NAT2_UGT73C6_* were due to a *bona fide* activity of the RNA. Here, we substituted the start codons of ORFs 1 to 3 by a CCC codon encoding proline (Figure 5A). We identified four independent homozygous lines that overexpressed the non-peptide coding *NAT2_UGT73C6_* (*npcNAT2_UGT73C6_*) sequence at high levels (*OE-npcNAT2_UGT73C6_* lines 1 to 4, Supplemental Figure 8), and assessed the impact of the mutated *NAT2_UGT73C6_* sequence on the identified phenotype. Interestingly, we found that in all the *OE-npcNAT2_UGT73C6_* lines, the rosette area was increased in comparison with the Col-0 wild-type controls (Figure 5B). Another important finding was that the values obtained in rosette area measurements with these lines correlated closely with those obtained in lines overexpressing the unmodified (wild-type) *NAT2_UGT73C6_* sequence (Figure 5B), two of which (*OE-NAT2_UGT73C6_* lines 5 and 6) were generated *de novo* for this comparison.

Overall, these data indicate that, although the peptides can be clearly detected when transiently overexpressed as GFP C-terminal fusions in *N. benthamiana*, their overexpression in *A. thaliana* does not lead to an increase in rosette size. On the other hand, as with the wild-type *NAT2_UGT73C6_*, overexpression of the *npcNAT2_UGT73C6_* variant leads to an increase in rosette area, suggesting that the observed phenotype depends on properties related to the RNA molecule itself.

Above, we reported that the downregulation of *NATs_UGT73C6_* transcripts leads to a reduction in the number of mesophyll and epidermal cells in leaves (Figure 3C), while *NATs_UGT73C6_* overexpression generates an increase of the rosette area. In view of this scenario, we decided to quantify cell size in leaf 6 of 35-day-old wild-type plants and plants overexpressing *NAT2_UGT73C6_* (*OE-NAT2_UGT73C6_-6*) and the *npcNAT2_UGT73C6_* variant (*OE-npcNAT2_UGT73C6_-3*), respectively. For this purpose, we harvested leaves from representative plants and performed cell-size measurements of abaxial epidermal and mesophyll cells in the tip and in the bottom regions. Interestingly, however, mesophyll and epidermal cells of either the *NAT2_UGT73C6_-* or the *npcNAT2_UGT73C6_*-overexpressing incr1e8ase in area compared to the control (Figure 5D), suggesting that the observed differences in leaf size are, at least in part, also due to cell enlargement. Estimations of the total cell number based on the mean cell size (Figure 5D) and determined leaf area (Figure 5C) revealed a significant increase of both cell types in the *NAT2_UGT73C6_* and the *npcNAT2_UGT73C6_* overexpressing lines (Figure 5E). Thus, on the one hand, these data confirmed the earlier finding (Figures 3 and 4) showing that the presence and expression level of the *NATs_UGT73C6_* modulates cell number and, consequently leaf and rosette size. On the other hand, we obtained some evidence that also an increase in cell size contributes to the *NATs_UGT73C6_*-mediated phenotype.

### Ectopic alteration of *NATs_UGT73C6_* expression does not affect the transcript levels of the *UGT73C6* sense gene or that of its closely related homologue *UGT73C5*

In an earlier report it has been proposed that in the UGT multigene family, lncNATs can modulate the expression of the sense gene and also of closely related genes by acting in *trans* (Wang et al., 2006). Hence, we wanted to understand whether such a scenario also applies to the studied *NATs_UGT73C6_*, the corresponding sense gene *UGT73C6* and its closest homologue *UGT73C5*.

The availability of the transgenic *A. thaliana* lines with altered *NATs_UGT73C6_* levels allowed us to quantify the expression of *UGT73C6* and *UGT73C5* in these lines. Surprisingly, we observed that the expression of *UGT73C6* and *UGT73C5* was not decreased in homozygous plants overexpressing *NAT1_UGT73C6_* or *NAT2_UGT73C6_* (Figure 6A and Supplemental Figure 9). Consistently, we found that the expression of *UGT73C6* and *UGT73C5* was also not altered in lines with reduced *NAT_UGT73C6_* levels (Figure 6A and Supplemental Figure 9).

Altogether, these results revealed that, at least in the natural host *A. thaliana*, ectopic changes in the expression of the *NATs_UGT73C6_* transcripts do not affect their complementary mRNAs and that the resulting leaf phenotype is not a consequence of significant changes in the expression of *UGT73C6* and *UGT73C5*.

### Brassinosteroid treatment does not consistently affect the levels of *NATs_UGT73C6_* transcripts, and brassinosteroid marker genes are not changed in lines with altered *NATs_UGT73C6_* expression

Our data suggested that *NATs_UGT73C6_* have no direct effect on the expression of *UGT73C6*, an important player in brassinosteroid (BR) homeostasis (Husar et al., 2011). To gain further insights into a possible ro20le for the *NATs_UGT73C6_* in this context, we decided to analyze the gene responses to externally applied brassinosteroid. To this end, we treated 8-day-old wild-type seedlings with 1 µM 24-epiBL, collected samples at different incubation times and determined gene expression levels by RT-qPCR. Consistent with a previous report (Husar et al., 2011), we found that expression of *UGT73C6* and of *UGT73C5* was not significantly altered (Figure 6B). Moreover, we observed that the treatment also had little effect on *NATs_UGT73C6_* expression, with a slight increase in expression after 3 hours, which, however, was not maintained after longer incubation periods (Figure 6B). Control genes analyzed to monitor the experimental conditions behaved as expected: *i.e.*, expression of *PHYTOCHROME P450 90B1* (*DWF4*) and *BRASSINOSTEROID-6-OXIDASE 2* (*BR-6-OX2*), genes repressed by BR application, was significantly decreased, while expression of *PHYB ACTIVATION- TAGGED SUPRESSOR 1* (*BAS1*), a catabolic gene induced under the same conditions (Tanaka et al., 2005), was significantly increased (Figure 6B). Further expression analyses at early time points revealed that neither *NATs_UGT73C6_* nor *UGT73C6* nor *UGT73C5* levels were affected after short periods of BR-treatment (Supplemental Figure 10A).

Thus, under the given conditions, the expression of *NATs_UGT73C6_* was not consistently altered in response to BR treatment.

Next, we considered it important to understand, whether the phenotypes, which resulted from altered *NATs_UGT73C6_* levels were caused by changes in the content of active brassinosteroid. To address this, we determined, in lines with altered *NATs_UGT73C6_* levels, the expression of *ROTUNDIFOLIA3* (*ROT3*), a BR biosynthetic gene, and *EXPANSIN 8* (*EXP8*), a BR-responsive gene of the expansin family (Goda et al., 2002). The results from these analyses showed that expression of *ROT3* was not affected when *NATs_UGT73C6_* were either overexpressed or downregulated, and that *EXP8* expression was slightly reduced in the line with decreased *NATs_UGT73C6_* levels (Figure 6C). In the lines overexpressing *UGT73C5*, *ROT3* expression was slightly increased and *EXP8* levels were significantly reduced (Supplemental Figure 10B), which was consistent with the described role of UGT73C5 in BR inactivation (Poppenberger et al., 2005).

These results suggest that active BR levels are not significantly changed in lines with altered *NATs_UGT73C6_* expression and, consequently, are not associated with the observed phenotype.

### Alteration of *NATs_UGT73C6_* expression leads to changes in *GRFs* transcript levels

Our results suggested that changes in *NATs_UGT73C6_* expression affect leaf and rosette size mainly by altering the number of mesophyll and epidermal cells. In the experiments with our promoter-GUS reporter lines we observed that *NAT2_UGT73C6_* was expressed at the bottom region of young leaves (Supplemental Figures 1A and 11A) and that its expression decreased during development. Interestingly, this pattern matches that of the members of the *GRF* family of transcriptional regulators, which are strongly expressed in actively growing and developing tissues (Rodriguez et al., 2010). GRFs and the transcriptional coactivator ANGUSTIFOLIA 3/GRF-INTERACTIONG FACTOR 1 (AN3/GIF1), which physically interacts with six of the nine proteins encoded by the *GRF* family members, are key regulators of leaf growth and achieve this by modulating cell proliferation (Vercruysse et al., 2020). The progression of leaf development is further controlled by microRNA396 (miR396), which regulates the expression of seven of the nine *GRFs* at the post-transcriptional level (Jones-Rhoades and Bartel, 2004; Liu et al., 2009; Rodriguez et al., 2010). In accord with this, we identified a potential binding site for miR396 in the *NATs_UGT73C6_* sequence (Supplemental Figure 11B), supporting the idea that *NATs_UGT73C6_* transcripts may have the potential to act as endogenous target mimics (Franco-Zorrilla et al., 2007). In this way, alteration in *NATs_UGT73C6_* expression could cause changes in *GRFs* levels.

To address this hypothesis, we decided to determine the transcript levels of *GRFs* and *GIF1* in the bottom and tip regions of young developing leaves of transgenic lines with altered *NATs_UGT73C6_* expression. For this purpose, we dissected leaf 3 from 12-day-old Arabidopsis seedlings grown on plates. We selected this time point because it has been previously reported that both cell proliferation and expansion continue to occur in the bottom and the tip regions of the leaf, respectively (Andriankaja et al., 2012). Analyses of the obtained RT-qPCR data showed that the levels of several GRF mRNAs, including those of *GRF4*, *6* and *9*, were increased in the line overexpressing *NAT2_UGT73C6_* (Figure 6D). In addition, we detected a statistically significant increase of *GIF1* levels in both leaf regions (Figure 6D). On the other hand, in *_amiR_NATs_UGT73C6_-10*, the line with reduced *NATs_UGT73C6_* levels, we detected a statistically significant reduction in *GRF6* transcripts at the leaf bottom (Figure 6E). Notably, *GRF6* was found to be the only analyzed *GRF* whose expression levels were changed in the lines where *NAT2_UGT73C6_* transcripts were overexpressed or downregulated. Interestingly, we observed here a positive correlation: *i.e.*, in comparison to the appropriate controls, the *GRF6* level was found to be increased when *NAT2_UGT73C6_* was overexpressed, while it was decreased when *NATs_UGT73C6_* were downregulated (Figures 6D and 6E). These data suggest that the level of this particular *GRF* is critically related to the observed phenotypes.

Altogether, these results suggest that *NATs_UGT73C6_* modulate *GIF1* and *GRFs* levels, which in turn, affect cell proliferation leading to changes in leaf size and rosette area.

## Discussion

Here we present the functional characterization of two lncNATs transcribed from a gene overlapping the *UGT73C6* gene from *A. thaliana*. The function of the enzymes encoded by *UGT73C6* and its closely related homologue *UGT73C5* have been elucidated earlier (Poppenberger et al., 2005; Husar et al., 2011) but nothing was known about the expression pattern and potential regulatory functions of the *NATs_UGT73C6_* transcripts. First, and by using reporter lines and qPCR analysis, we confirmed that both *NATs_UGT73C6_* are expressed and that the promoters are active in different organs. Furthermore, we demonstrated that the expression of the *NATs_UGT73C6_* is developmentally controlled and that it takes place independently of the presence of the sense gene in *cis*. Similar observations have been made previously with other lncNATs in plants (Jabnoune et al., 2013; Fedak et al., 2016) reinforcing the idea that the identified lncNATs are functionally relevant and do not correspond to transcriptional ‛background noise’ resulting from sense gene expression.

Several of the earlier characterized lncNATs from plants act in *trans* on their targets when ectopically overexpressed. These examples include *asHSFB2a*, which controls gametophyte development (Wunderlich et al., 2014) and *FLORE,* which modulates the onset of flowering (Henriques et al., 2017) in *Arabidopsis*, and *cis-NATPHO1;2*, that regulates phosphate homeostasis in rice (Jabnoune et al., 2013). *Trans*-acting lncNATs are known to be relatively stable and to recruit other factors to regulate their targets. We found the half-life of *NATs_UGT73C6_* to be comparable to that of a canonical mRNA, a feature indeed that suggests a *trans*-acting function.

In our studies we demonstrate that changes in *NATs_UGT73C6_* expression levels, which were achieved by post-transcriptional downregulation by amiRNAs or overexpression under the control of the strong *35S* promoter, result in mild but consistent phenotypes in which the size of individual leaves and, consequently, rosette area was affected (Figures 2 and 4). In lines with a *NATs_UGT73C6_* downregulation close to 50 percent the rosette area was reduced by almost 20 percent in comparison to controls where the *NATs_UGT73C6_* transcript levels were unaffected. Interestingly, reductions in size were observed with all leaves and were maintained also in fully developed leaves indicating that this phenotype is not a consequence of delayed growth. On the other hand, overexpression of either of the *NATs_UGT73C6_* transcripts resulted in plants with increased rosette area. The same phenotype was detectable when full-length, non-peptide coding variants of *NAT2_UGT73C6_* were overexpressed while, in contrast, it was undetectable when partial-length variants with peptide-coding capacity were overexpressed. The later observations were important especially given the fact that both *NATs_UGT73C6_* contain several ORFs and that one of them encodes for a 103 amino acid long peptide when translated from the unspliced *NATs_UGT73C6_* variant. These results support the conclusion that the identified phenotype is due to the activity of the RNA molecules themselves, which thus function as *bona fide* lncRNAs.

Our analyses at the cellular level showed that in the line where *NATs_UGT73C6_* were downregulated there were no significant differences in mesophyll cell size and that the size of the abaxial epidermal cells was slightly increased. Accordingly, both observations could not explain the reduction in leaf size. However, when we estimated the total number of cells, we observed a clear decrease for both cell types, suggesting that cell proliferation is affected. On the other hand, in lines with increased expression of the wild-type *NAT2_UGT73C6_* or of the non-peptide coding *npcNAT2_UGT73C6_* variant the cell area of both cell types was found to be increased. While these size differences again did not explain the changes observed in leaf area, we obtained further data indicating that the cell number is also increased in the overexpression lines. Thus, in sum our data suggest that the main differences in leaf size that were observed in lines with altered *NATs_UGT73C6_* levels result from changes in cell proliferation.

As explained above, NATs may control the expression of a sense transcript in *cis* or in *trans*. Given the high sequence similarity of *UGT73C*-subfamily members, we originally assumed that the *NATs_UGT73C6_* would regulate the expression of *UGT73C6* and/or other closely related *UGTs*, as has been reported for *FLORE*, which represses multiple *CDFs (CYCLING DOF FACTORS)* in *cis* and *trans* (Henriques et al., 2017). However, our data surprisingly show that overexpression of *NATs_UGT73C6_* did not reduce the levels of either *UGT73C6* or its closest homologue *UGT73C5*. These results are congruent with those reported by Deforges and coworkers who obtained transgenic lines with high overexpression of *cis-*NATs in which sense gene expression remained unaffected (Deforges et al., 2019). On the other hand, our data contrast with the “Yin-Yang” mechanism proposed for the pair formed by *HSFB2a* and *asHSFB2a*, in which overexpression of either member of the pair leads to knockdown of the other (Wunderlich et al., 2014). Consistently, post-transcriptional downregulation of *NATs_UGT73C6_* by amiRNAs was found to have no effect on *UGT73C6* or *UGT73C5* levels, indicating that the *NATs_UGT73C6_* do not modulate the expression of these genes or the levels of their transcripts in *trans*. Unfortunately, the *NATs_UGT73C6_* transcription initiation sites locate within the coding sequence of *UGT73C6* making it difficult, if not impossible, to study their effect on *UGT73C6* expression in *cis*. Therefore, we cannot exclude potential *cis* effects of *NATs_UGT73C6_* transcription on *UGT73C6* expression. However, our data strongly suggest that the *NATs_UGT73C6_* are not involved in brassinosteroid homeostasis. This opinion is supported by the fact that the expression of *UGT73C6* or *UGT73C5* remained essentially unaffected even in the presence of strong changes in *NATs_UGT73C6_* levels. Furthermore, no link between *NATs_UGT73C6_* and hormone levels could be established.

By contrast, our data suggest that the observed, distinct phenotypes are a direct consequence of the activities of the *NATs_UGT73C6_* transcripts. Based on this premise, we decided to analyze the expression of other potential targets. Considering (i) the observed changes in the number of cells in response to the alteration of the expression of *NATs_UGT73C6_* (Figures 3 and 5), (ii) the pattern of *NAT2_UGT73C6_* promotor activity, which is mainly detected at the bottom region of young developing leaves (Supplemental Figure 11A), and, (iii) the presence of a potential binding site with partial complementarity to miR396 in the *NATs_UGT73C6_* sequence (Supplemental Figure 11B), we focused our analysis on the members of the *GRF* family, which, together with GIF1, are crucial for cell number determination in leaves (Vercruysse et al., 2020). Members of the *GIF* and *GRF* gene families are predominantly expressed at the bottom region of young leaves (Rodriguez et al., 2010), and the levels of most of the *GRF* family members are regulated by miR396 (Jones-Rhoades and Bartel, 2004). Overexpression of *GIF1* and of several *GRFs* results in plants with larger organs due to increased cell proliferation (Lee et al., 2009; Vercruysse et al., 2020). *gif1* mutants, on the other hand, show the opposite phenotype, *i.e.*, plants with smaller and narrower leaves as a result of reduced cell number (Kim and Kende, 2004). Overexpression of miR396 leads to post-transcriptional downregulation of *GRFs* containing its target site and to plants with narrow leaves due to cell number reduction (Liu et al., 2009). Conversely, plants transformed with miR396- resistant variants of *GRFs* display a higher cell number and larger leaves (Rodriguez et al., 2010; Debernardi et al., 2014).

miRNA activity can be modulated by endogenous target mimicry, a mechanism in which non-target transcripts may bind complementary miRNAs. Due to mismatches at particular positions in the binding site no further processing occurs by the RNAi machinery, i.e., RNA-induced silencing complexes (RISC). Thus, active miRNAs are sequestered, which in turns, leads to protection and increased levels of the miRNÁs direct targets. The first and best-characterized example of this mechanism is the long noncoding RNA *INDUCED BY PHOSPHATE STARVATION1* (*IPS1*) (Franco-Zorrilla et al., 2007), the expression of which is induced after phosphate starvation. By binding and sequestering miR399, *IPS1* protects and promotes the translation of the mRNA of *PHO*, which encodes a E2 ubiquitin conjugase-related protein.

When *NAT2_UGT73C6_* is overexpressed, we detected a significant increase in the levels of *GRF6*, as well as of *GRF4* and *GRF9* transcripts at the bottom and tip regions of the leaves, respectively (Figure 6D). These data indeed suggest that the *NATs_UGT73C6_* act as target mimics that sequester miR396 and thus protect miR-targeted *GRF* mRNAs from RISC activity. Such a scenario fits with data obtained in transgenic plants expressing an artificial target mimic directed against miR396, in which a two-fold increased level of miR396-regulated *GRFs* was observed (Debernardi et al., 2012). Our results show that the effect for *GRF4* and *GRF9* is more pronounced in the tip region (Figure 6D). This is particularly relevant considering that miR396 expression in young leaves follows a gradient along the longitudinal axis with higher levels in this region (Debernardi et al., 2012). Thus, ectopic overexpression of *NAT2_UGT73C6_* could cause an increase of *GRFs* transcript levels in this leaf region by protecting them against miR396 activity. We detected a reduction in *GRF6* transcripts at the bottom leaf region in the line where *NATs_UGT73C6_* were downregulated (Figure 6E). Conversely, we observed an increase in the same region in the line that overexpressed *NAT2_UGT73C6_* (Figure 6D), suggesting that the presence of the *NATs_UGT73C6_* is important for maintaining wild-type *GRF6* mRNA levels. It is interesting to note that both, *GRF6* and *GRF5* lack the miR396 binding site. Despite this fact, *GRF6* expression is reduced in plants overexpressing the miRNA suggesting a feed-forward effect of the other *GRFs* on its levels (Rodriguez et al., 2010). In addition, *GIF1* expression was also found to be significantly increased in both leaf regions of the *NAT2_UGT73C6_* overexpressing line (Figure 6D).

The GRF-GIF duo is known to activate their own transcription through a positive feedback regulatory loop (Vercruyssen et al., 2014). Accordingly, increases in *GRFs* due to the target mimic protective action of *NATs_UGT73C6_* could lead to an increment of *GIF1* transcripts in the *NAT2_UGT73C6_* overexpression line. Simultaneous expression of high levels of *GRF3* and *GIF1* transcripts results in a synergistic increase in leaf cell number and additive effects on gene expression (Debernardi et al., 2014), suggesting that the phenotype in the overexpression lines may be enhanced by the concurrent increased levels of *GRFs* and *GIF1*.

Interestingly, the potential binding site of miR396 in the *NATs_UGT73C6_* transcripts does not follow the canonical requirements of complementarity at the 5’ end (Supplemental Figure 11B). In fact, it is at the critical value of allowed mismatches (≤5) that is the hallmark of functionally verified miRNA target sites in plants (Liu et al., 2014). A weak *NATs_UGT73C6_*- *miR396* interaction may be relevant to avoid strong competition for miR396 binding, which could lead to detrimental effects. This can be relevant considering that lncNATs are usually involved in a fine-tuning of gene expression (Kindgren et al., 2018).

The fact that the *NATs_UGT73C6_* are present in young leaves and that their levels drop when *GRFs* are no longer expressed supports the idea that *NATs_UGT73C6_* activities are particularly relevant to protect the corresponding mRNAs from miR396 action at early developmental stages. The relatively long half-life of *NATs_UGT73C6_* and the presence of splicing variants support the idea of a cytoplasmic localization of the transcripts, which in turn, is in agreement with their potential role as miRNA target mimics (Noh et al., 2018). While the observed miR396 binding site is present in almost all transcript variants that were tested during these studies, we observed an increase in rosette area exclusively when we overexpressed the full-length *NATs_UGT73C6_* molecules (Figure 4 and Figure 5B). Different scenarios can explain this intriguing finding. On the one hand, it is conceivable that the function(s) of the *NATs_UGT73C6_* are only enabled by a complex structure that is formed by the full-length RNAs, containing, for example, the complete 3’-end. Such a situation has been described for *HIDDEN TREASURE 1 (HID1)* (Wang et al., 2014), a lncRNA containing four main stem-loops of which two are essential for its function, the regulation of *PHYTOCHROME-INTERACTING FACTOR 3* (*PIF3*) gene expression. On the other hand, these data may be indicative of a protein, which for example, may be required for exposure of the miR396-binding site and that binds exclusively to the full-length *NATs_UGT73C6_* molecules.

It was previously reported that *TWISTED LEAF* (*TL*), an endogenous lncRNA from rice, influences leaf blade shape by modulating the expression of the *R2R3 MYB* transcription factor gene with which it overlaps (Liu et al., 2018). To our knowledge, our work represents the first report of lncNATs involved in regulation of leaf size, adding an additional layer of complexity to the process of leaf growth.

## Methods

### Plant material and growth conditions

*Arabidopsis thaliana* ecotype Col-0 was used as wild-type background. All generated transgenic lines and the knockout mutant *ugt73c6ko* (SAIL_525_H07) were in the same genetic background (Col-0). Seeds were surfaced-sterilized, stratified for 3 days in water (4°C, in the dark) and plated on ½ plates containing Murashige and Skoog (MS) basal medium supplemented with 1% (w/v) sucrose. *A. thaliana* seedlings were grown in a growth chamber under long-day conditions (16 h of light/ 8 h of dark) at 22°C/ 20°C. To evaluate phenotypes on adult plants 10-day-old seedlings were transferred to individual 8 cm diameter pots filled with a steam-sterilized soil and maintained in growth chambers (Percival Scientific) under long-day conditions and controlled temperature (22°C/ 20°C) and humidity (65-70%), or in the greenhouse.

Seeds of the *ugt73c6_ko_* mutant (SAIL_525_H07) were obtained from the Nottingham Arabidopsis Stock Centre (NASC). Homozygous plants were identified by PCR genotyping, and transcript levels were determined by quantitative RT-PCR (RT-qPCR). **Transgenic lines generation**

For the generation of *NAT1_UGT73C6_* and *NAT2_UGT73C6_* overexpressing plants the sequences reported in the TAIR10 database were amplified by PCR from genomic DNA extracted from *A. thaliana* plants. The same template was used for the amplification of promoter regions of *NAT1_UGT73C6_* (2054 bp) and *NAT2_UGT73C6_* (2580 bp) starting immediately upstream of the respective reported transcription initiation site. The genomic DNA was also used to amplify the promoter regions of *UGT73C6* (bases -1687 to +4 considering the translation initiation site) and *UGT73C5* (759 bp upstream of the translation initiation site) based on previous publications (Poppenberger et al., 2003; Husar et al., 2011). Amplicons starting with a non-template CACC sequence were directionally cloned into the pENTR^TM^ /D-TOPO vector (Life Technologies, USA).

The pENTR clone containing the genomic *NAT2_UGT73C6_* sequence was used as template for start codon(s) removal by site-directed mutagenesis. Overlapping primers in which ATG sequences were replaced by the CCC trinucleotide were designed for each start codon and utilized in PCR reactions with AccuPrime Pfx DNA polymerase (Thermo Scientific, USA) following the manufactureŕs instructions. The plasmid containing the insert with the desired mutations, introduced by repeated rounds of mutagenesis, was used as donor for gateway cloning.

Potential protein-coding capacity from both *NATs_UGT73C6_* was analyzed using the ExPASy translator tool online (https://web.expasy.org/translate/), and protein domains were analyzed using the webtool of the Pfam database (www.pfam.xfam.org) and PROSITE (www.prosite.expasy.org). Primers were designed to amplify the corresponding ORFs without the triplet encoding the stop codon and including or excluding the *NAT2_UGT73C6_* upstream sequence. Amplification was performed using the pENTR clone containing the genomic *NAT2_UGT73C6_* sequence as template and amplicons were directionally cloned into the pENTR^TM^ /D-TOPO vector (Life Technologies, USA).

Sequence identity and absence of undesired mutations was confirmed by sequencing (Eurofins Genomics, Germany) the inserts that were present in all selected pENTR clones. Inserts from the recombinant pENTR plasmids were transferred to destination vectors *via* gateway cloning by using the LR clonase^TM^ II Enzyme mix (Invitrogen, USA) and by following the manufactureŕs instructions. pB7WG2, pBGWFS7, and pB7FWG2 vectors (Karimi et al., 2002) were used to generate overexpression, promoter-reporter and GFP C-terminal fusion lines, respectively.

amiRNA candidates were selected using the P-SAMS amiRNA Designer tool (http://p-sams.carringtonlab.org/) (Carbonell et al., 2014). Constructs and clones for plant transformation were generated according to the protocol provided by the authors. Destination vectors pMDC32B-AtMIR390a-B/c and pMDC123SB-AtMIR390a-B/c (Carbonell et al., 2014) were used, which allowed plant selection with hygromycin or with the glufosinate-ammonium herbicide (BASTA^®^, Bayer), respectively.

Presence and identity of the cloned products in the pDEST vectors were determined by colony PCR and restriction pattern analysis. Binary plasmids were introduced by electroporation into the *Agrobacterium* strain GV3101, and suspensions from transformed bacteria were used for *A. thaliana* transformation by the floral-dip method (Clough and Bent, 1998). Transgenic plants were selected based on Mendelian segregation of the resistance gene. Expression levels of the transgenes were analyzed by northern blot or RT-qRT-PCR. At least three independent homozygous lines were selected for analysis.

### RT-qRT-PCR

Total RNA was extracted from Arabidopsis seedlings and rosette leaves using the peqGOLD Plant RNA kit (PeqLab) according to the manufactureŕs instructions. Genomic DNA potentially present after purification was removed using double-stranded DNase (dsDNase, Thermo Fisher Scientific). Total RNA was reverse transcribed into cDNA with SuperScript II reverse transcriptase (Thermo Fisher Scientific) using oligo dT or gene-specific primers. cDNAs were amplified by PCR using Phusion or Dream-Taq (Thermo Fisher Scientific), or by RT-qRT-PCR with a QuantStudio5 Real-Time instrument (Thermo Fisher Scientific) using FAST SYBR^TM^ Green qPCR Master Mix (Thermo Fisher Scientific). Relative expression was calculated and normalized to internal reference genes [*protein phosphatase 2A subunit* (*PP2AA3*, *At1g13320*) or *glyceraldehyde-3-phosphate dehydrogenase C*-2 (*GAPC2*, *At1g13440*) (Czechowski et al., 2005)]. For each experiment, at least three biological replicates and two to three technical replicates for each sample were performed, and experiments were repeated at least twice. ΔC_t_ values were calculated as C_t (reference gene)_-C_t (gene of interest)_, where higher C_t_ values indicate higher expression of the gene of interest. ΔΔC_t_ values represent the mean difference in ΔC_t_ between treatment and control samples (positive and negative ΔΔC_t_ values indicates induction or repression due to the treatment, respectively).

Primers used in this study are listed in Supplemental Data Set 1.

### Reporter localization analysis

*GUS* staining was performed according to standard protocols. Briefly, seedlings of homozygous transgenic lines carrying the *GUS*-reporter constructs were incubated overnight at 37°C in *GUS* staining solution [0.5 mM K_4_Fe(CN)_6_, 0.5 mM K_3_Fe(CN)_6_, 0.1% Triton X100, 0.5 mg/mL X-Gluc in DMF, 50 mM NaH_2_PO_4_ pH7.0]. Chlorophyll was removed by incubation in aqueous solutions containing increasing ethanol concentrations (20, 35, 50 and 70% (v/v) final concentration). Older leaves, flowers and siliques were fixed for 30 min with a solution containing 90% (v/v) absolute ethanol and 10% acetic acid and further incubated for 1 h in 90% (v/v) ethanol. The final preparations were stored in 70% (v/v) ethanol. Pictures were taken with a SMZ1270® stereomicroscope (NIKON).

### Brassinosteroid treatment

The experiments were performed as previously reported (Husar et al., 2011). Briefly, Col-0 seedlings were grown on vertical plates containing ½ MS media for 5 days, transferred to sterile flasks containing liquid ½ MS media and incubated for 2 days in continuous light conditions with agitation. 24-epi-brassinolide (epiBL, Sigma Aldrich®) dissolved in DMSO was added to a 1 µM final concentration and seedlings were further incubated under the described conditions. Control seedlings were supplemented with the same volume of DMSO. Samples were collected at different time points (0, 10, 30 and 60 min, 2, 3 and 24 h), snap frozen in liquid nitrogen and conserved at -80°C until analyzed.

### RNA half-life determination

RNA half-lives were determined as previously described (Fedak et al., 2016). Col-0 seedlings were grown during 11 days on vertical plates containing ½ MS medium and then transferred to a 6-well-plate containing incubation buffer (1 mM PIPES pH 6.25, 1 mM trisodium citrate, 1 mM KCl, 15 mM sucrose). After 30 min of incubation, cordycepin (3’ deoxyadenosine) was added to a final concentration of 150 mg/L and internalized using vacuum (2 cycles of 5 min each). Seedlings were collected at regular time-points every 15 min (0, 15, 30, 45, 60, 75, 90, 105 and 120 min) and frozen in liquid nitrogen. Total RNA was isolated using the peqGOLD Plant RNA Kit (PeqLab), reverse transcribed, and gene expression analyzed by RT-qRT-PCR. Short- and long-lived transcripts *EXPANSIN-LIKE1 (Expansin L1*) and *Eukaryotic translation initiation factor 4A1* (*EIF4A1A*) were used as unstable and stable controls, respectively. C_t_-values were normalized by the C_t_-value at time point 0 [C_t(n)_ = (ln(C_t_/C_t(0)_)*(-10)] and plotted as degradation curves. Slopes of the curves were used to calculate RNA half-lives [t_1/2_ = (ln_2_)/slope].

### Rosette and leaf area measurements

Complete plant pictures were recorded with a digital camera (Sony Sony® DSLR). For rosette area determination digital images corresponding to 20 to 30 plants per line were analyzed with the Easy Leaf Area program (Easlon and Bloom, 2014) and the green area quantified. According to the instructions, a red calibrator of a fixed area was included in the pictures. Plants for dissection were grown under similar conditions, individual leaves were cut, and their area was measured using ImageJ (Schneider et al., 2012).

### Microscopic analysis and cell size determination

Leaf material for microscopic analysis was prepared according to Nelissen and collaborators (Nelissen et al., 2013) with modifications. Leaf 6 from selected 35-day-old plants were incubated in fixation solution (90% absolute ethanol, 10% acetic acid) for 2 h followed by overnight incubation in 90% ethanol. Samples were placed in 70% ethanol for 30 minutes and stored in lactic acid. Two hours before microscopy the samples were transferred to Hoyer medium (80 g chloral hydrate in 30 ml water) for clearing. Leaves were cut in four sections following the longitudinal axis and pictures from mesophyll and abaxial epidermal cells were taken from the tip and the bottom regions by differential interference contrast (DIC) microscopy using an Apotome.2 microscope (Zeiss) and an Axiocam ERc 5s microscope camera (Zeiss). Ten or more pictures were taken from each sample (leaves from the two plants closer to the median in the rosette area determination). Complete cell borders were drawn manually either digital on the photo using the Vectornator program (https://www.vectornator.io) or analoguosly after printing the photos on transparent foil, scanned for digitalization and processed with the processing package Fiji (https://fiji.sc/) (Schindelin et al., 2012) to obtain individual cell area. To quantify only fully developed epidermal pavement cells an inferior limit size of 2000 µm^2^ was used.

### Cell number estimation

Total palisade mesophyll and abaxial epidermal cell number in leaf 6 was estimated by dividing the determined area of the individual leaves by the average cell area for each cellular type. For epidermal cells, an upper limit of 15000 µm^2^ was fixed to exclude abnormally big cells (less than 4% for all lines except for *OE-NAT2_UGT73C6_-6*, in which they represent the 5.7%).

### Statistical analysis

Data were tested either with unpaired two-tailed Student’s *t*-Test or with one-way ANOVA and Tukeýs multiple comparison test using the GraphPad Prism 7 software (GraphPad Software, San Diego, CA, USA). Graphs were generated using the same software.

## Supplemental Data

Supplemental Figure 1. Histochemical localization of GUS activity at different developmental stages in the Pro*NAT1_UGT73C6_:GUS* and Pro*NAT2_UGT73C6_:GUS* transgenic lines.

Supplemental Figure 2. Identification of *NAT2_UGT73C6_* splice variants.

Supplemental Figure 3. Gene expression analysis and rosette area quantification in the *ugt73c6_ko_* line.

Supplemental Figure 4. Identification and characterization of transgenic lines with reduced *NATs_UGT73C6_* expression.

Supplemental Figure 5. Expression analysis in transgenic lines transformed with the *NATs_UGT73C6_* sequences and leaf growth kinematics.

Supplemental Figure 6. Detection of small peptides encoded by *NAT2_UGT73C6_* after transient expression in *N. benthamiana*.

Supplemental Figure 7. Expression analysis and quantification of rosette area in *A. thaliana* plants overexpressing the small peptides encoded by *NAT2_UGT73C6_* as C-terminal GFP-fusions.

Supplemental Figure 8. Expression analysis in transgenic lines transformed with the wild-type *NAT2_UGT73C6_* sequence or with the non-peptide coding *NAT2_UGT73C6_* variant.

Supplemental Figure 9. U*G*T73C5 levels in lines with altered *NATs_UGT73C6_* expression.

Supplemental Figure 10. Early response to brassinosteroid treatment and expression of brassinosteroid marker genes.

Supplemental Figure 11. Histochemical localization of GUS activity in a young leaf of the Pro*NAT2_UGT73C6_:GUS* transgenic line and potential miRNA396 binding site in the *NATs_UGT73C6_* sequence.

## ACKNOWLEDGMENTS

We thank Alberto Carbonell (Instituto de Biología Molecular y Celular de Plantas, Spain) for the design of the amiGUS constructs and James Carrington (Donald Danforth Plant Science Center, USA) for providing vectors for amiRNAs expression. We acknowledge Brigitte Poppenberger (Technical University Munich, Germany) for seeds of the *OE- UCT73C5* line. We thank Silvestre Marillonet and Sabine Rosahl (Leibniz Institute of Plant Biochemistry, Germany) for providing the p19 clone, and for the conceptual input and the comments on the manuscript, respectively. We thank Gary Sawers for reading the manuscript. We acknowledge Kristin Eismann and Michael André Fritz for technical assistance and initial data, respectively.

This work was funded by grants of the Deutsche Forschungsgemeinschaft (DFG) [RTG1591 (projects C2 and C3)], the European Union [ERASMUS MUNDUS Action 2 program “BRAVE”], the State Research Priority Program Saxony-Anhalt Fkz [ZS/2016/06/79740] and core funding from the state of Saxony-Anhalt and the Federal Republic of Germany to the Leibniz Institute of Plant Biochemistry (IPB). S.K.M. was funded by a BRAVE scholarship, by IPB and by projects C2 and C3 of the RTG1591. S.E. was funded by project C2 of the RTG1591 and by IPB. A.J. was funded by project C3 of the RTG1591. S.G.Z. was funded by the Priority Program and IPB.

## AUTHOR CONTRIBUTIONS

S.G.Z. designed the experiments and wrote the manuscript. S.K.M., M.H., S.E., A.J., T.d.V., K.B. and S.G.Z. generated the constructs and the transgenic lines, performed the experiments, and analyzed the results. S.A. and S.E.B. contributed with conceptual input and comments.

All authors read and approved the final article.

## Parsed Citations

Andriankaja, M., Dhondt, S., De Bodt, S., Vanhaeren, H., Coppens, F., De Milde, L., Muhlenbock, P., Skirycz, A., Gonzalez, N., Beemster, G.T., and Inze, D. (2012). Exit fromproliferation during leaf development in Arabidopsis thaliana: a not-so-gradual process. Dev Cell 22, 64–78.

Ariel, F., Romero-Barrios, N., Jegu, T., Benhamed, M., and Crespi, M. (2015). Battles and hijacks: noncoding transcription in plants. Trends in plant science 20, 362–371.

Ben Amor, B., Wirth, S., Merchan, F., Laporte, P., d’Aubenton-Carafa, Y., Hirsch, J., Maizel, A., Mallory, A., Lucas, A., Deragon, J.M., Vaucheret, H., Thermes, C., and Crespi, M. (2009). Novel long non-protein coding RNAs involved in Arabidopsis differentiation and stress responses. Genome research 19, 57–69.

Borsani, O., Zhu, J., Verslues, P.E., Sunkar, R., and Zhu, J.K. (2005). Endogenous siRNAs derived froma pair of natural cis-antisense transcripts regulate salt tolerance in Arabidopsis. Cell 123, 1279–1291.

Caputi, L., Malnoy, M., Goremykin, V., Nikiforova, S., and Martens, S. (2012). Agenome-wide phylogenetic reconstruction of family 1 UDP-glycosyltransferases revealed the expansion of the family during the adaptation of plants to life on land. The Plant journal : for cell and molecular biology 69, 1030–1042.

Carbonell, A., Takeda, A., Fahlgren, N., Johnson, S.C., Cuperus, J.T., and Carrington, J.C. (2014). New generation of artificial MicroRNA and synthetic trans-acting small interfering RNAvectors for efficient gene silencing in Arabidopsis. Plant physiology 165, 15–29.

Clark, M.B., Johnston, R.L., Inostroza-Ponta, M., Fox, A.H., Fortini, E., Moscato, P., Dinger, M.E., and Mattick, J.S. (2012). Genome-wide analysis of long noncoding RNAstability. Genome research 22, 885–898.

Clough, S.J., and Bent, A.F. (1998). Floral dip: a simplified method for Agrobacterium-mediated transformation of Arabidopsis thaliana. The Plant journal : for cell and molecular biology 16, 735–743.

Csorba, T., Questa, J.I., Sun, Q., and Dean, C. (2014). Antisense COOLAIR mediates the coordinated switching of chromatin states at FLC during vernalization. Proceedings of the National Academy of Sciences of the United States of America 111, 16160–16165.

Cui, J., Luan, Y., Jiang, N., Bao, H., and Meng, J. (2017). Comparative transcriptome analysis between resistant and susceptible tomato allows the identification of lncRNA16397 conferring resistance to Phytophthora infestans by co-expressing glutaredoxin. The Plant journal : for cell and molecular biology 89, 577–589.

Czechowski, T., Stitt, M., Altmann, T., Udvardi, M.K., and Scheible, W.R. (2005). Genome-wide identification and testing of superior reference genes for transcript normalization in Arabidopsis. Plant physiology 139, 5–17.

Debernardi, J.M., Rodriguez, R.E., Mecchia, M.A., and Palatnik, J.F. (2012). Functional specialization of the plant miR396 regulatory network through distinct microRNA-target interactions. PLoS Genet 8, e1002419.

Debernardi, J.M., Mecchia, M.A., Vercruyssen, L., Smaczniak, C., Kaufmann, K., Inze, D., Rodriguez, R.E., and Palatnik, J.F. (2014). Post-transcriptional control of GRF transcription factors by microRNA miR396 and GIF co-activator affects leaf size and longevity. The Plant journal : for cell and molecular biology 79, 413–426.

Deforges, J., Reis, R.S., Jacquet, P., Sheppard, S., Gadekar, V.P., Hart-Smith, G., Tanzer, A., Hofacker, I.L., Iseli, C., Xenarios, I., and Poirier, Y. (2019). Control of Cognate Sense mRNA Translation by cis-Natural Antisense RNAs. Plant physiology 180, 305–322.

Easlon, H.M., and Bloom, A.J. (2014). Easy Leaf Area: Automated digital image analysis for rapid and accurate measurement of leaf area. Appl Plant Sci 2.

Fedak, H., Palusinska, M., Krzyczmonik, K., Brzezniak, L., Yatusevich, R., Pietras, Z., Kaczanowski, S., and Swiezewski, S. (2016). Control of seed dormancy in Arabidopsis by a cis-acting noncoding antisense transcript. Proceedings of the National Academy of Sciences of the United States of America 113, E7846–E7855.

Franco-Zorrilla, J.M., Valli, A., Todesco, M., Mateos, I., Puga, M.I., Rubio-Somoza, I., Leyva, A., Weigel, D., Garcia, J.A., and Paz-Ares, J. (2007). Target mimicry provides a new mechanism for regulation of microRNA activity. Nat Genet 39, 1033–1037.

Goda, H., Shimada, Y., Asami, T., Fujioka, S., and Yoshida, S. (2002). Microarray analysis of brassinosteroid-regulated genes in Arabidopsis. Plant physiology 130, 1319–1334.

Gonzalez, N., Vanhaeren, H., and Inze, D. (2012). Leaf size control: complex coordination of cell division and expansion. Trends in plant science 17, 332–340.

Henriques, R., Wang, H., Liu, J., Boix, M., Huang, L.F., and Chua, N.H. (2017). The antiphasic regulatory module comprising CDF5 and its antisense RNAFLORE links the circadian clock to photoperiodic flowering. The New phytologist 216, 854–867.

Husar, S., Berthiller, F., Fujioka, S., Rozhon, W., Khan, M., Kalaivanan, F., Elias, L., Higgins, G.S., Li, Y., Schuhmacher, R., Krska, R., Seto, H., Vaistij, F.E., Bowles, D., and Poppenberger, B. (2011). Overexpression of the UGT73C6 alters brassinosteroid glucoside formation in Arabidopsis thaliana. BMC plant biology 11, 51.

Jabnoune, M., Secco, D., Lecampion, C., Robaglia, C., Shu, Q., and Poirier, Y. (2013). Arice cis-natural antisense RNAacts as a translational enhancer for its cognate mRNA and contributes to phosphate homeostasis and plant fitness. The Plant cell 25, 4166–4182.

Jones-Rhoades, M.W., and Bartel, D.P. (2004). Computational identification of plant microRNAs and their targets, including a stress-induced miRNA. Mol Cell 14, 787–799.

Jones, P., Messner, B., Nakajima, J., Schaffner, A.R., and Saito, K. (2003). UGT73C6 and UGT78D1, glycosyltransferases involved in flavonol glycoside biosynthesis in Arabidopsis thaliana. The Journal of biological chemistry 278, 43910–43918.

Kalve, S., Fotschki, J., Beeckman, T., Vissenberg, K., and Beemster, G.T. (2014). Three-dimensional patterns of cell division and expansion throughout the development of Arabidopsis thaliana leaves. J Exp Bot 65, 6385–6397.

Karimi, M., Inze, D., and Depicker, A. (2002). GATEWAY vectors for Agrobacterium-mediated plant transformation. Trends in plant science 7, 193–195.

Kim, J.H., and Kende, H. (2004). Atranscriptional coactivator, AtGIF1, is involved in regulating leaf growth and morphology in Arabidopsis. Proceedings of the National Academy of Sciences of the United States of America 101, 13374–13379.

Kindgren, P., Ard, R., Ivanov, M., and Marquardt, S. (2018). Transcriptional read-through of the long non-coding RNASVALKA governs plant cold acclimation. Nat Commun 9, 4561.

Lapidot, M., and Pilpel, Y. (2006). Genome-wide natural antisense transcription: coupling its regulation to its different regulatory mechanisms. EMBO reports 7, 1216–1222.

Lee, B.H., Ko, J.H., Lee, S., Lee, Y., Pak, J.H., and Kim, J.H. (2009). The Arabidopsis GRF-INTERACTING FACTOR gene family performs an overlapping function in determining organ size as well as multiple developmental properties. Plant physiology 151, 655–668.

Li, Y., Baldauf, S., Lim, E.K., and Bowles, D.J. (2001). Phylogenetic analysis of the UDP-glycosyltransferase multigene family of Arabidopsis thaliana. The Journal of biological chemistry 276, 4338–4343.

Lin, S., Zhang, L., Luo, W., and Zhang, X. (2015). Characteristics of Antisense Transcript Promoters and the Regulation of Their Activity. Int J Mol Sci 17.

Liu, D., Song, Y., Chen, Z., and Yu, D. (2009). Ectopic expression of miR396 suppresses GRF target gene expression and alters leaf growth in Arabidopsis. Physiol Plant 136, 223–236.

Liu, Q., Wang, F., and Axtell, M.J. (2014). Analysis of complementarity requirements for plant microRNA targeting using a Nicotiana benthamiana quantitative transient assay. The Plant cell 26, 741–753.

Liu, X., Li, D., Zhang, D., Yin, D., Zhao, Y., Ji, C., Zhao, X., Li, X., He, Q., Chen, R., Hu, S., and Zhu, L. (2018). Anovel antisense long noncoding RNA, TWISTED LEAF, maintains leaf blade flattening by regulating its associated sense R2R3-MYB gene in rice. The New phytologist 218, 774–788.

Mercer, T.R., Dinger, M.E., and Mattick, J.S. (2009). Long non-coding RNAs: insights into functions. Nat Rev Genet 10, 155–159.

Nelissen, H., Rymen, B., Coppens, F., Dhondt, S., Fiorani, F., and Beemster, G.T. (2013). Kinematic analysis of cell division in leaves of mono- and dicotyledonous species: a basis for understanding growth and developing refined molecular sampling strategies. Methods in molecular biology 959, 247–264.

Noh, J.H., Kim, K.M., McClusky, W.G., Abdelmohsen, K., and Gorospe, M. (2018). Cytoplasmic functions of long noncoding RNAs. Wiley interdisciplinary reviews. RNA 9, e1471.

Poppenberger, B., Berthiller, F., Lucyshyn, D., Sieberer, T., Schuhmacher, R., Krska, R., Kuchler, K., Glossl, J., Luschnig, C., and Adam, G. (2003). Detoxification of the Fusarium mycotoxin deoxynivalenol by a UDP-glucosyltransferase from Arabidopsis thaliana. The Journal of biological chemistry 278, 47905–47914.

Poppenberger, B., Fujioka, S., Soeno, K., George, G.L., Vaistij, F.E., Hiranuma, S., Seto, H., Takatsuto, S., Adam, G., Yoshida, S., and Bowles, D. (2005). The UGT73C5 of Arabidopsis thaliana glucosylates brassinosteroids. Proceedings of the National Academy of Sciences of the United States of America 102, 15253–15258.

Rinn, J.L., and Chang, H.Y. (2012). Genome regulation by long noncoding RNAs. Annual review of biochemistry 81, 145–166.

Rodriguez, R.E., Mecchia, M.A., Debernardi, J.M., Schommer, C., Weigel, D., and Palatnik, J.F. (2010). Control of cell proliferation in Arabidopsis thaliana by microRNA miR396. Development 137, 103–112.

Rosa, S., Duncan, S., and Dean, C. (2016). Mutually exclusive sense-antisense transcription at FLC facilitates environmentally induced gene repression. Nat Commun 7, 13031.

Ross, J., Li, Y., Lim, E., and Bowles, D.J. (2001). Higher plant glycosyltransferases. Genome Biol 2, REVIEWS3004.

Schindelin, J., Arganda-Carreras, I., Frise, E., Kaynig, V., Longair, M., Pietzsch, T., Preibisch, S., Rueden, C., Saalfeld, S., Schmid, B., Tinevez, J.Y., White, D.J., Hartenstein, V., Eliceiri, K., Tomancak, P., and Cardona, A. (2012). Fiji: an open-source platformfor biological- image analysis. Nat Methods 9, 676–682.

Schneider, C.A., Rasband, W.S., and Eliceiri, K.W. (2012). NIH Image to ImageJ: 25 years of image analysis. Nat Methods 9, 671–675.

Scholthof, H.B. (2006). The Tombusvirus-encoded P19: fromirrelevance to elegance. Nature reviews. Microbiology 4, 405–411.

Tanaka, K., Asami, T., Yoshida, S., Nakamura, Y., Matsuo, T., and Okamoto, S. (2005). Brassinosteroid homeostasis in Arabidopsis is ensured by feedback expressions of multiple genes involved in its metabolism. Plant physiology 138, 1117–1125.

Vercruysse, J., Baekelandt, A., Gonzalez, N., and Inze, D. (2020). Molecular networks regulating cell division during Arabidopsis leaf growth. J Exp Bot 71, 2365–2378.

Vercruyssen, L., Verkest, A., Gonzalez, N., Heyndrickx, K.S., Eeckhout, D., Han, S.K., Jegu, T., Archacki, R., Van Leene, J., Andriankaja, M., De Bodt, S., Abeel, T., Coppens, F., Dhondt, S., De Milde, L., Vermeersch, M., Maleux, K., Gevaert, K., Jerzmanowski, A., Benhamed, M., Wagner, D., Vandepoele, K., De Jaeger, G., and Inze, D. (2014). ANGUSTIFOLIA3 binds to SWI/SNF chromatin remodeling complexes to regulate transcription during Arabidopsis leaf development. The Plant cell 26, 210–229.

Wang, H., Chua, N.H., and Wang, X.J. (2006). Prediction of trans-antisense transcripts in Arabidopsis thaliana. Genome Biol 7, R92.

Wang, H.V., and Chekanova, J.A. (2017). Long Noncoding RNAs in Plants. Adv Exp Med Biol 1008, 133–154.

Wang, Y., Fan, X., Lin, F., He, G., Terzaghi, W., Zhu, D., and Deng, X.W. (2014). Arabidopsis noncoding RNA mediates control of photomorphogenesis by red light. Proceedings of the National Academy of Sciences of the United States of America 111, 10359–10364.

Wunderlich, M., Gross-Hardt, R., and Schoffl, F. (2014). Heat shock factor HSFB2a involved in gametophyte development of Arabidopsis thaliana and its expression is controlled by a heat-inducible long non-coding antisense RNA. Plant molecular biology 85, 541–550.

Yao, R.W., Wang, Y., and Chen, L.L. (2019). Cellular functions of long noncoding RNAs. Nat Cell Biol 21, 542–551.

Yu, Y., Zhang, Y., Chen, X., and Chen, Y. (2019). Plant Noncoding RNAs: Hidden Players in Development and Stress Responses. Annu Rev Cell Dev Biol 35, 407–431.

Zhang, X., Lii, Y., Wu, Z., Polishko, A., Zhang, H., Chinnusamy, V., Lonardi, S., Zhu, J.K., Liu, R., and Jin, H. (2013). Mechanisms of small RNAgeneration fromcis-NATs in response to environmental and developmental cues. Molecular plant 6, 704–715.

Zhao, X., Li, J., Lian, B., Gu, H., Li, Y., and Qi, Y. (2018). Global identification of Arabidopsis lncRNAs reveals the regulation of MAF4 by a natural antisense RNA. Nat Commun 9, 5056.

